# Selective modulation of cell surface proteins during vaccinia infection: implications for immune evasion strategies

**DOI:** 10.1101/2021.10.06.463320

**Authors:** Delphine M Depierreux, Arwen F Altenburg, Lior Soday, Alice Fletcher-Etherington, Robin Anthrobus, Brian J Ferguson, Michael P Weekes, Geoffrey L Smith

**Author notes:** Joint corresponding authors. Authors contributed equally to this study (Delphine M Depierreux, Arwen F Altenburg).

## Abstract

The interaction between immune cells and virus-infected targets involves multiple plasma membrane (PM) proteins. A systematic study of PM protein modulation by vaccinia virus (VACV), the paradigm of host regulation, has the potential to reveal not only novel viral immune evasion mechanisms, but also novel factors critical in host immunity. Here, >1000 PM proteins were quantified throughout VACV infection, revealing selective downregulation of known T and NK cell ligands including HLA-C, downregulation of cytokine receptors including IFNAR2, IL-6ST and IL-10RB, and rapid inhibition of expression of certain protocadherins and ephrins, candidate activating immune ligands. Downregulation of most PM proteins occurred via a proteasome-independent mechanism. Upregulated proteins included a decoy receptor for TRAIL. Twenty VACV-encoded PM proteins were identified, of which five were not recognised previously as such. Collectively, this dataset constitutes a valuable resource for future studies on antiviral immunity, host-pathogen interaction, poxvirus biology, vector-based vaccine design and oncolytic therapy.

## Introduction

Vaccinia virus (VACV) is a large, double-stranded (ds)DNA virus and is best known as the live vaccine used to eradicate smallpox (1). Since smallpox eradication in 1980, research with VACV has continued because it is an excellent model to study host-pathogen interactions. Furthermore, VACV was developed as an expression vector (2, 3) with utility as a vaccine against other infectious diseases (4-6) and as an oncolytic agent (7, 8). To optimise the design of VACV-based vaccines and oncolytic agents, it is important to develop a comprehensive understanding of the interactions between VACV-infected cells and the host immune system, and how VACV modulates these interactions. The study of virus-induced changes to cellular proteins has also led to several advances in understanding of host cell protein function. A recent example of this was our demonstration that histone deacetylase 4 (HDAC4) is degraded during VACV infection (9), is needed for the recruitment of STAT2 to the interferon (IFN)-stimulated response element during type I IFN-induced signalling and restricts the replication of VACV and herpes simplex virus type 1 (10).

VACV encodes many immunomodulatory proteins that function to evade or suppress the host immune response to infection (11). Intracellular immunomodulators may inhibit innate immune signalling pathways, the activity of IFN-stimulated gene (ISG) products, or block apoptosis (12, 13). Secreted proteins can bind and inhibit cytokines, chemokines, IFNs or complement factors. Additional immunomodulators function on the cell surface to influence recognition of the infected cell by the immune system. In general, these have been studied less extensively than secreted or intracellular proteins. Nonetheless, it was reported that the viral haemagglutinin (HA, protein A56) modulates interactions with natural killer (NK) cells (14), and A40 is a cell surface protein with a type II membrane topology, with limited amino acid similarity to C-type lectins and NK cell receptors (15). In addition to these integral membrane proteins, some VACV secreted proteins can also bind to the surface of infected or uninfected cells. Examples include the type I IFN-binding protein (B18 in VACV strain Western Reserve – WR) that binds to cell surface glycosaminoglycans (16, 17) and the M2 protein that binds to B7.1 and B7.2 to prevent T cell activation (18, 19). The vaccinia complement control protein C3 (VCP) and the K2 serine protease inhibitor (serpin 3, SPI-3) each bind to A56 (20), and vaccinia epidermal growth factor (VGF) binds to the epidermal growth factor receptor and promotes cell division (21) and affects virus spread (22). Other VACV proteins present on the infected cell surface are part of the outer envelope of the extracellular enveloped virus (EEV) (23).

In addition to VACV proteins expressed at the cell surface, there have been a few reports of changes to cellular plasma membrane (PM) proteins during infection. For instance, the abundance of different MHC class I haplotypes, a major immune ligand, have been reported to change during infection and this might influence recognition of infected cells by both CD8^+^ cytotoxic T lymphocytes and NK cells (24-27). However, changes in cell surface protein expression during VACV infection have not been addressed comprehensively or systematically.

This study used plasma membrane profiling (PMP) (28, 29) to provide a comprehensive analysis of temporal and quantitative changes in host and viral proteins at the surface of immortalised human foetal foreskin fibroblasts (HFFF-TERTs) following VACV infection. Using tandem mass-tag (TMT)-based proteomics of PM-enriched fractions, >1000 PM proteins were quantified, and of these, 142 were downregulated and 113 were upregulated during infection. Twenty VACV proteins were detected at the cell surface including C8 and F5, which were not known to be present at the PM. Modulation of the expression levels of PM proteins indicated several possible novel immune evasion strategies, including selective downregulation of HLA-C and the IFN-α receptor 2 (IFNAR2) and upregulation of an apoptosis decoy receptor for TRAIL. The use of a proteasome inhibitor and comparison with previous studies assessing whole cell protein expression during VACV infection (9) and protein stability (30) in HFFF-TERTs suggested that proteasomal degradation and host protein synthesis shut-off are not the major mechanisms by which PM proteins are downregulated during VACV infection, and that most PM proteins are likely upregulated through subcellular translocation and/or stabilisation at the PM. Finally, a comparative analysis with a dataset examining cell surface proteomic changes upon infection of HFFF-TERTs with human cytomegalovirus (HCMV) (31) identified possible common viral immune evasion strategies.

## Results

### Quantitative temporal analysis of the plasma membrane proteome during VACV infection

To measure how VACV infection changes the cell surface proteome, HFFF-TERTs were mock-treated or infected with VACV WR in biological duplicate. These cells were used previously in an investigation of the whole cell proteome during VACV infection (9) and the whole cell lysate (WCL) and PM proteomes during HCMV infection (29), thereby enabling direct comparisons with these datasets. Additionally, a single mock and an infected sample were treated with the proteasome inhibitor MG132 at 2 hours post-infection (hpi). Flow cytometry confirmed that >95% of the cells were infected (**Figure S1A-B**). Multiplexed TMT and triple-stage mass spectrometry (MS3) were used to quantify the relative abundance of PM proteins at 1.5, 6, 12 and 18 hpi (**Figure 1A**).

**Figure 1.**
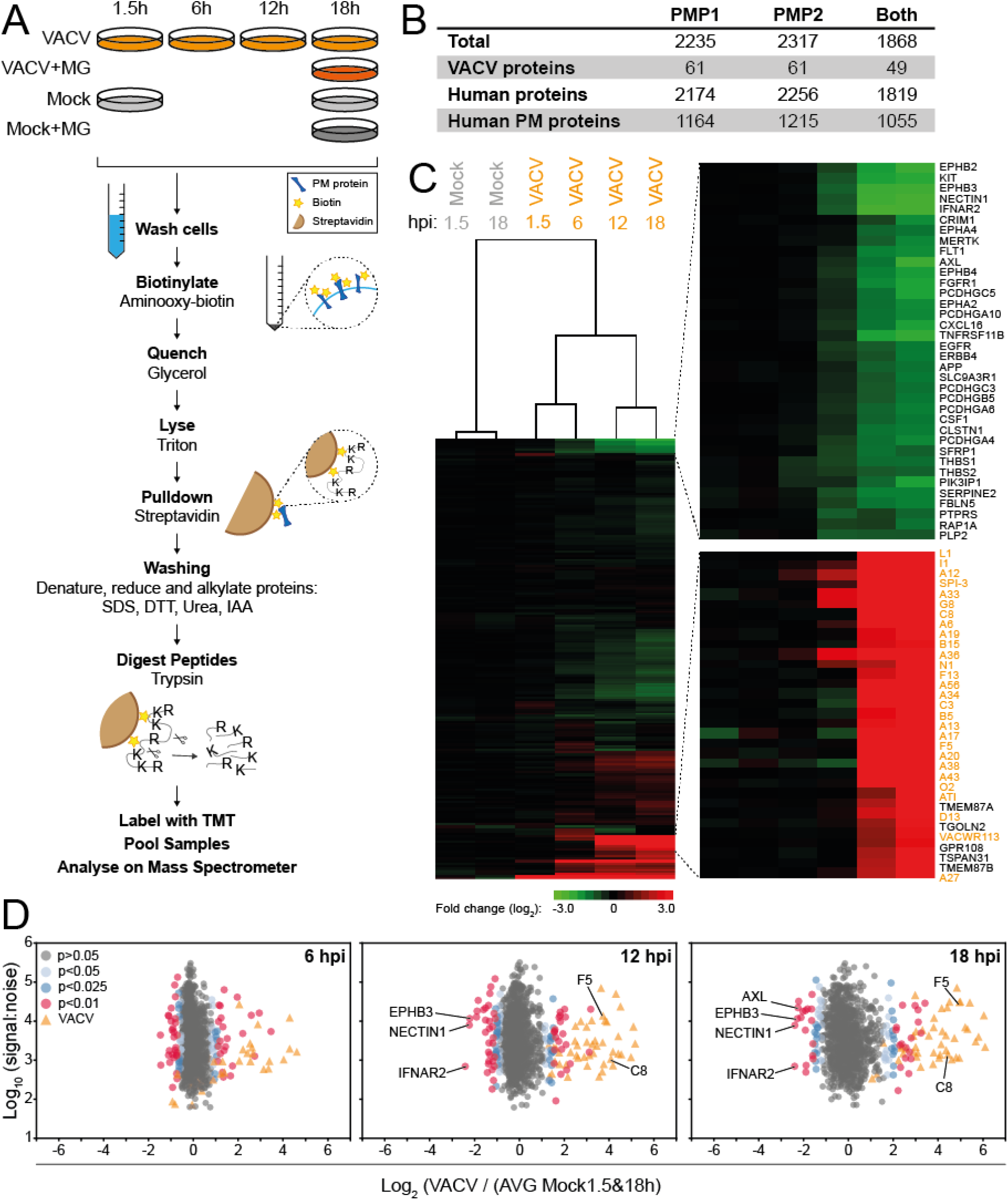
Quantitative temporal analysis of the plasma membrane proteome during VACV infection. (**A**) Schematic of the experimental workflow. HFFF-TERTs were mock-treated or infected with VACV at MOI 5 for the indicated time-points (**Fig. S1A-B**). At 2 hpi MG132 was added to a mock and an infected sample (‘+MG’). Samples were generated in biological duplicate (PMP1, PMP2). (**B**) Number of proteins quantified in the PMP replicates. ‘Human PM proteins’ represents the number of proteins annotated with relevant GO terms (PM, ‘cell surface’ [CS], ‘extracellular’ [XC] and ‘short GO’ [ShG, 4-part term containing ‘integral to membrane’, ‘intrinsic to membrane’, ‘membrane part’, ‘cell part’ or a 5-part term additionally containing ‘membrane’]). (**C**) Hierarchical cluster analysis showing the fold change of all VACV and human proteins quantified in both replicates compared to mock (average 1.5 and 18 h, **Fig. S1C**). Selected sections are shown enlarged and VACV proteins are indicated in orange. (**D**) Scatter plots of all VACV and human PM proteins quantified in both repeats at 6, 12 or 18 hpi. Selected human PM proteins were annotated. P-values were estimated using significance A with Benjamini-Hochberg correction for multiple hypothesis testing (32).

Human proteins were filtered for gene ontology (GO) annotations related to PM expression. Overall, 49 VACV proteins and 1055 human PM proteins were quantified in both experiments (**Figure 1B**). Mock-infected samples presented negligible variation in the abundance of any given protein over the course of the experiment (**Figure S1C**). VACV-infection induced selective changes in the expression of PM proteins, with the greatest fold-change (FC) occurring mostly late during infection (**Figure 1C-D**). This was reflected by separate clustering of mock samples, and samples harvested early (1.5 & 6 hpi) or late (12 & 18 hpi) after VACV infection (**Figure 1C**). All data are shown in **Table S1**, in which the worksheet ‘‘Plotter’’ enables interactive generation of temporal graphs of the expression of each human or viral proteins quantified.

### Selective changes in human cell surface protein expression following VACV infection

Two sets of criteria were defined to determine which human PM proteins showed altered levels during VACV infection (**Figure 2A, Table S2A-D**). First, ‘sensitive’ criteria included proteins quantified in either or both PMP replicates showing >2-fold change (FC) at any time-point during infection. Second, ‘stringent’ criteria included only proteins detected in both PMP replicates showing >2 FC with a p-value <0.05 (Benjamini-Hochberg corrected one-way ANOVA). Both criteria indicated that VACV infection selectively alters the abundance of a small fraction (∼1%) of human PM proteins detected in this study (**Figure 2A, Table S2A-D**). Sensitive criteria were used for subsequent analyses and proteins identified by stringent criteria are shown where appropriate in supplementary tables.

**Figure 2.**
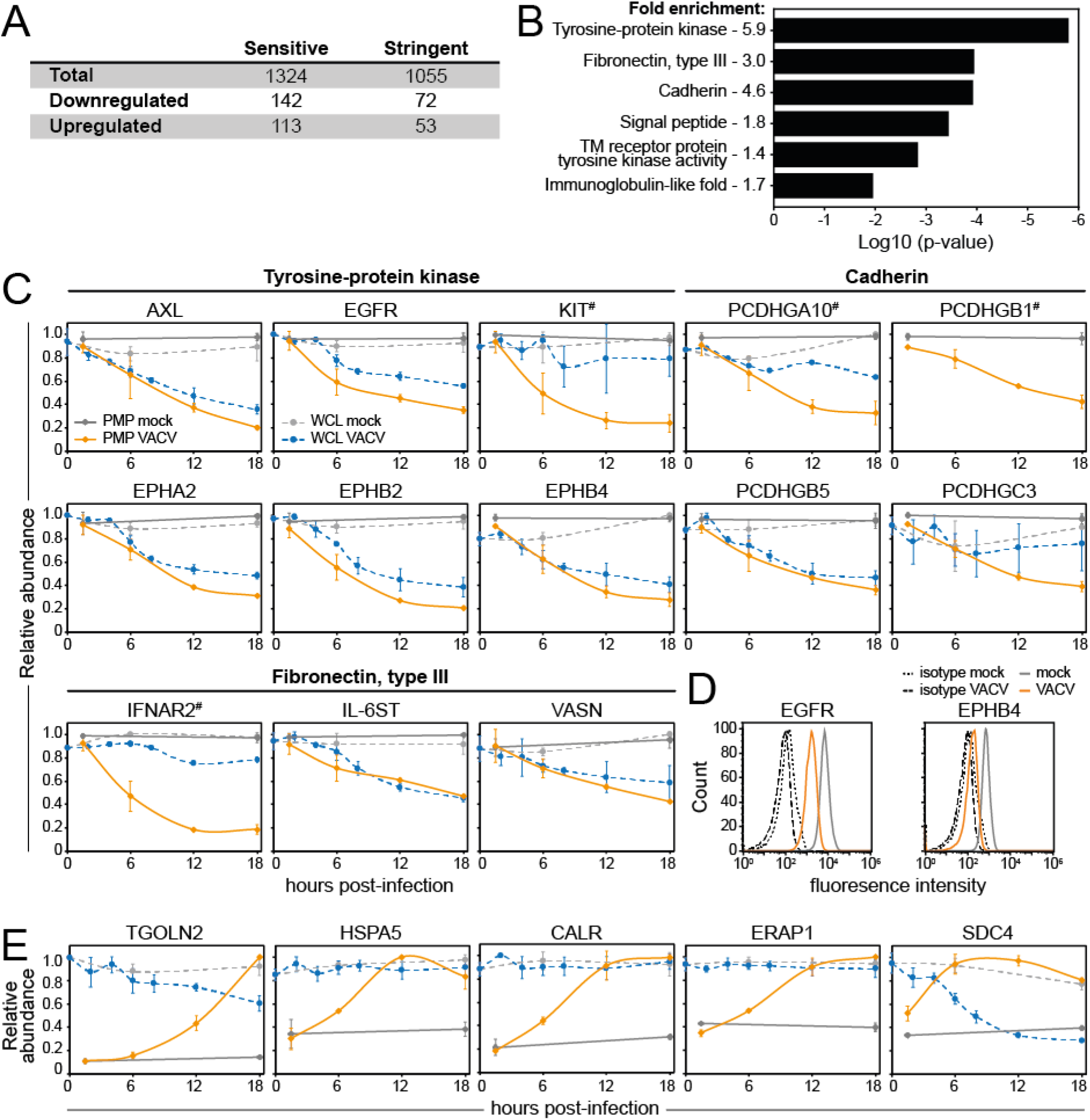
Selective modulation of host proteins at the cell surface during VACV infection. (**A**) Human host PM modulated according to the sensitive and stringent criteria (**Table S2A-D**). (**B**) DAVID functional enrichment of 142 proteins detected in either repeat and downregulated >2-fold. A background of all quantified human PM proteins was used. Representative terms are shown for each cluster with a Benjamini-Hochberg-corrected p-value <0.05 (**Table S2E**). (**C**) Temporal profiles of selected downregulated proteins, in which the fold-change for downregulation was in each case significant at p<0.05 (Benjamini-Hochberg-corrected one-way ANOVA, **Table S2A/C**). (**D**) Downregulation of EGFR/EPHB4 during VACV infection was confirmed by flow cytometry at 15 hpi with VACV (MOI 5). Results are representative of 3 independent experiments. (**E**) Temporal profiles of selected upregulated proteins, all with p <0.05 (Benjamini-Hochberg-corrected one-way ANOVA, **Table S2B/D**). Data are represented as mean ± SD (PMP n=2; WCL (9) n=3, # WCL n<3).

Of the 1055 human PM proteins quantified in both PMP replicates, 142 and 113 proteins were down- or upregulated, respectively (**Figure 2A, Table S2A-B**). The Database for Annotation, Visualization and Integrated Discovery (DAVID) (33, 34) identified six functional clusters that were enriched within the group of downregulated human PM proteins (**Figure 2B, Table S2E**). This included protocadherins and several clusters associated with receptor tyrosine kinases (RTKs), which contain immunoglobulin domains (growth factor receptor families), fibronectin type III domains (ephrin family), or a combination of the two (TAM family) (35). Temporal profiles provide insight into the kinetics of downregulation from the cell surface (this dataset) and as determined by WCL proteomics of VACV-infected cells (9) (**Figure 2C**). Downregulation of ephrin B4 (EPHB4) and epidermal growth factor receptor (EGFR) was confirmed by flow cytometry (**Figure 2D**).

DAVID functional enrichment analysis for the group of upregulated human PM proteins resulted in a single significantly enriched cluster: ‘Protein processing in the endoplasmic reticulum (ER)’ (**Table S2F**). The most highly upregulated proteins included many ER, Golgi and lysosomal proteins, such as trans-Golgi network integral membrane protein (TGOLN)2, heat shock protein (HSP)A5, calreticulin (CALR) and ER aminopeptidase (ERAP)1 (**Figure 2E**).

Interestingly, several host surface proteins involved with the cytokine response were modulated during VACV infection. For example, the interleukin-6 receptor subunit β (IL-6ST), interferon α/β receptor 2 (IFNAR2), mast/stem cell growth factor receptor Kit (KIT) (**Figure 2C**) and interleukin-10 receptor subunit β (IL-10RB, **Table S1**) were substantially downregulated from the PM during VACV infection. Conversely, PM expression of several proteins involved in the suppression of the cytokine response, including CALR, ERAP1 and syndecan-4 (SDC4), were upregulated (**Figure 2E**). Overall, PM expression modulation of these proteins may indicate novel strategies by which VACV manipulates the cytokine environment to enhance immune evasion, replication or spread.

### Selective modulation of cell surface immune ligands during VACV infection

NK and T cells are essential components of the antiviral immune response. Their activation status is determined by the integration of inhibitory and activating signals emanating from receptors engaging with ligands expressed by (infected) target cells. Interestingly, several of these immune ligands showed altered PM expression during VACV infection.

Major histocompatibility complex class I (MHC-I, human leukocyte antigen class I (HLA) in humans) molecules are important activators and regulators of NK and T cells. Due to the polymorphic nature of classical HLA-I (HLA-A, -B, -C), their peptides may easily be miss-assigned after detection by mass spectrometry. Therefore, only peptides corresponding uniquely to a single HLA-I type were included in this analysis. Interestingly, HLA-A and HLA-B were modestly downregulated whilst HLA-C was substantially downregulated (**Figure 3A**). This selective modulation was observed at both the cell surface and whole cell level and was further confirmed by flow cytometry in two different cell lines (**Figure 3B-C**). Given that all HLA-C subtypes are ligands for killer-cell immunoglobulin-like receptors (KIRs) expressed by NK cells, and less than 50% of the HLA-A/B subtypes can bind KIRs (36), these data suggest selective modulation of the NK cell response during VACV infection.

**Figure 3.**
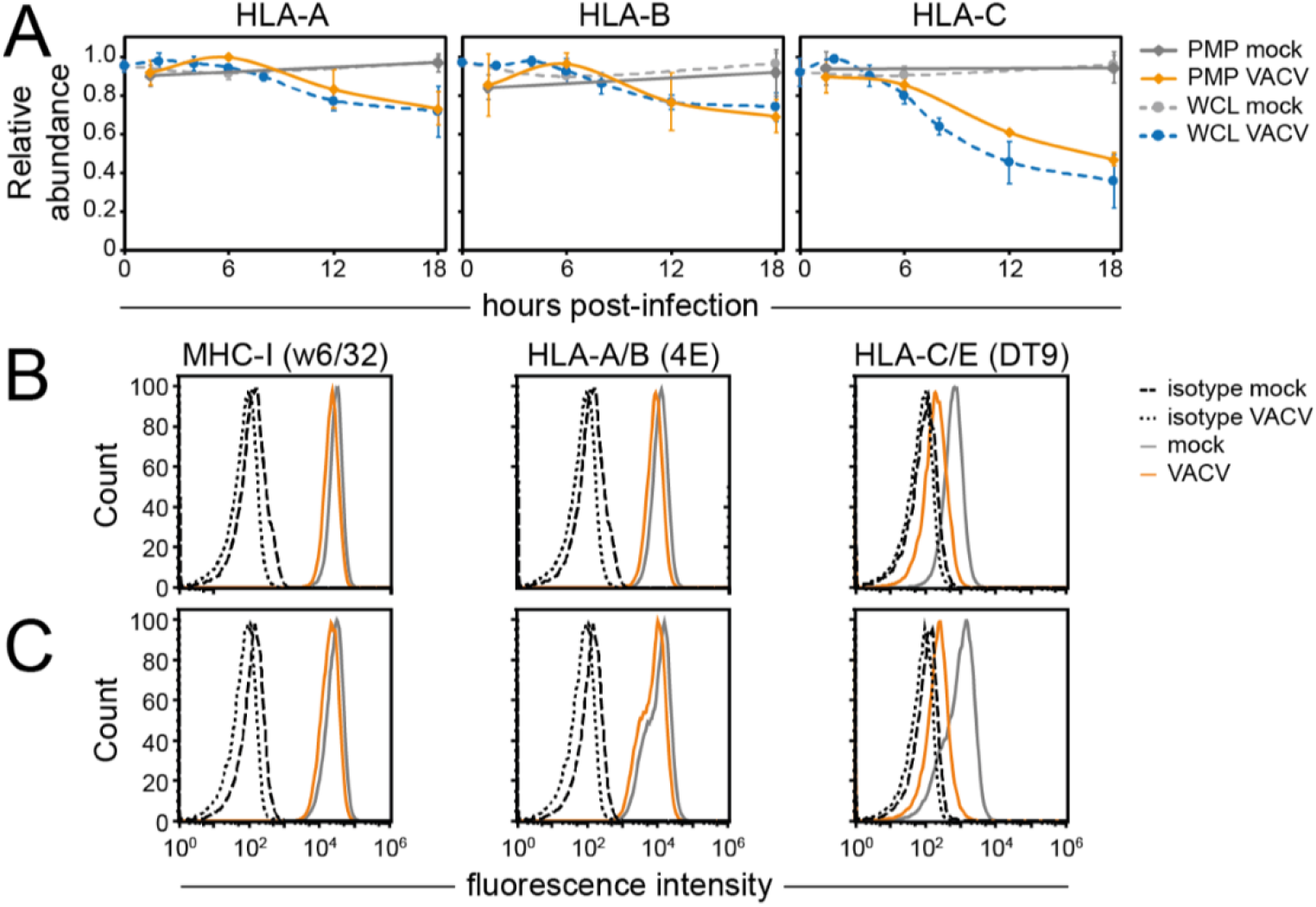
Selective downregulation of HLA-C from the PM during VACV infection. (**A**) Temporal profiles were generated only using peptides belonging uniquely to each of the indicated HLA-I heavy chains. Data are represented as mean ± SD (PMP n=2; WCL (9) n=3, # WCL n<3). (**B-C**) Cell surface downregulation of selected proteins during VACV infection (MOI 5) was confirmed by flow cytometry in HFFF-TERTs (**B**) or HeLa cells (**C**) at 15-18hpi. Results are representative of at least 2 independent experiments.

Enhanced PM expression of stress molecules such as NK activating ligands MHC class I polypeptide related sequence (MIC)A/B, UL-16-binding proteins (ULBPS) and B7-H6 represents a conserved cellular response to stress, including viral infection (37-39). However, the PM expression level of these proteins remained largely unchanged during VACV infection, which was confirmed by flow cytometry (**Figure 4A**). This may represent a new VACV strategy to evade immune recognition.

**Figure 4.**
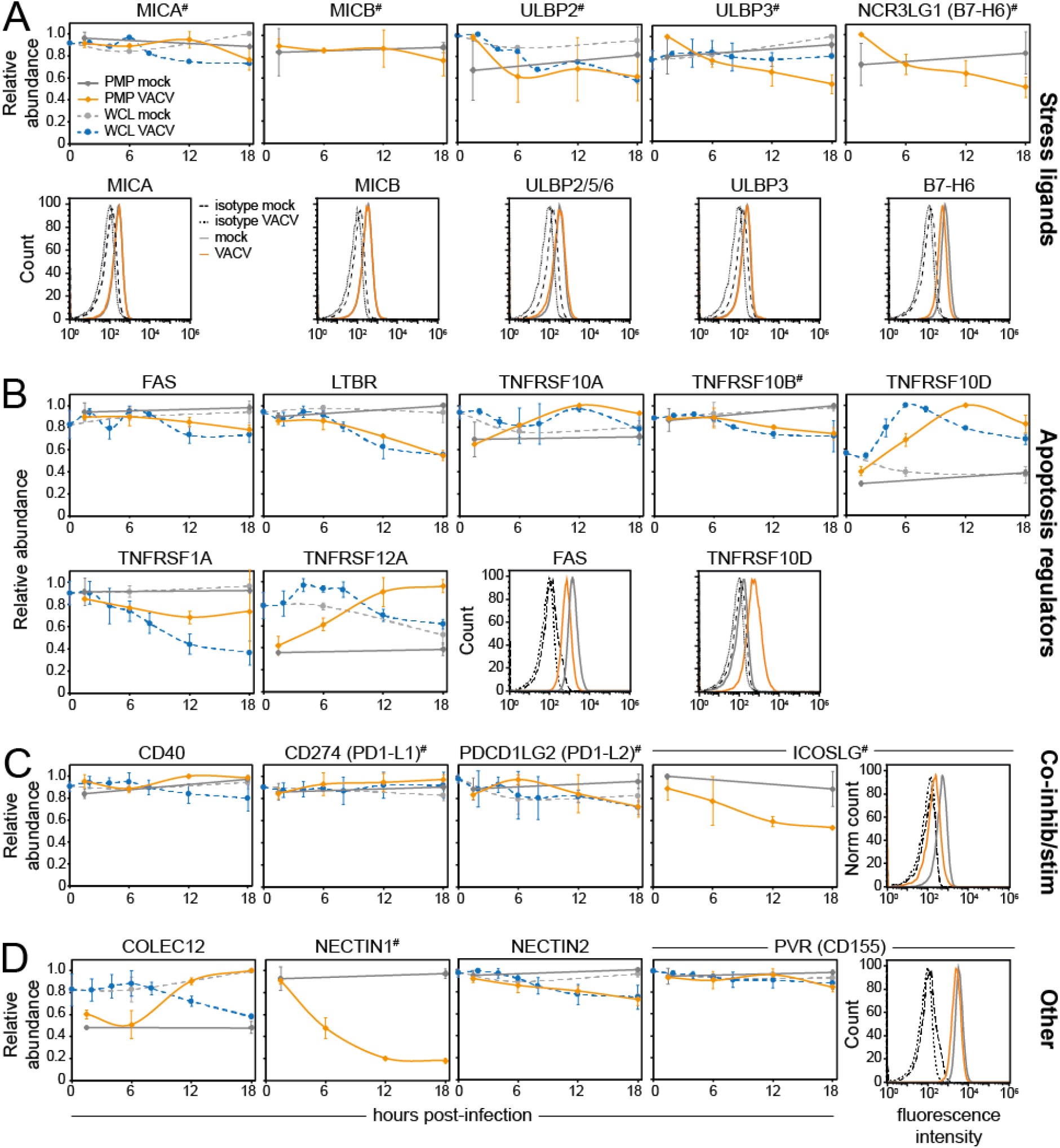
Cell surface expression of immune cell ligands is selectively modulated during VACV infection. Temporal profiles showing the cell surface and whole-cell expression levels of (**A**) stress ligands for NKG2D receptor, (**B**) apoptosis regulators, (**C**) co-inhibitory/stimulatory ligands and (**D**) other immune ligands. Data are represented as mean ± SD (PMP n=2; WCL (9) n=3, # WCL n<3). Cell surface downregulation of selected proteins during VACV infection was confirmed by flow cytometry in HeLa cells (stress ligands & TNFRSF10D), HFFF-TERTs (FAS) or DOHH2 cells (ICOSLG) at 15-18 hpi. Results are representative of at least two independent experiments.

Selective modulation of regulators of lymphocyte-mediated apoptosis was observed during VACV infection. Downregulation of lymphotoxin-β receptor (LTBR) and tumour necrosis factor receptor superfamily member (TNFRSF) 1A (TNFR1), as well as upregulation of TNFRSF10D (TRAIL-R4), a decoy receptor for TNF-related apoptosis-inducing ligand (TRAIL), may lead to decreased sensitivity to apoptosis (**Figure 4B**). Upregulation of TNFRSF10D was confirmed by flow cytometry (**Figure 4B**). Conversely, upregulation of apoptosis inducer and NF-κB activator TNFRSF12A (Fn14) (**Figure 4B**) may sensitise the infected cell to lymphocyte-induced apoptosis. Expression levels of other surface proteins involved in apoptosis regulation, including FAS and TNFRSF10A/-B, remained largely unchanged (**Figure 4B, S2**).

Immune checkpoints are activating and inhibitory pathways that regulate the delicate balance between lymphocyte activation and maintenance of self-tolerance. During VACV infection, the levels of inhibitory checkpoint molecules programmed cell death ligand (PD-L)1, PD-L2 and B7-H3 were stable (**Figure 4C, S2**). The temporal profile of activating checkpoint molecule CD40 also remained mostly unchanged (**Figure 4C**). In contrast, repulsive guidance molecule B (RGMB), which has both stimulatory and inhibitory functions, was downregulated from the PM (**Figure S2E**). Additionally, inducible costimulatory ligand (ICOSLG) was downregulated from the cell surface, which was confirmed by flow cytometry (**Figure 4C**).

Other noteworthy changes in the surface proteome during VACV infection include the upregulation of collectin-1 (COLEC12), a ligand for the inhibitory NK receptor paired immunoglobulin-like type 2 receptor α (PILRα, **Figure S2F**). Furthermore, NECTIN-1, a ligand for the CD96 receptor with both inhibitory and stimulatory properties, was substantially downregulated. Other related proteins such as poliovirus receptor (PVR or CD155), NECTIN-2 and NECTIN-3 remained unchanged, suggesting that NECTIN-1 is targeted selectively by VACV. These changes may represent previously unrecognised NK cell immunomodulatory strategies employed by VACV. Conversely, modulation of surface expression levels of activating ligand vimentin (VIM, **Figure S2E**), may reflect the host antiviral response and enhance sensitivity to NK cell killing.

Lymphocytes rely on adhesion molecules to make contact with surrounding cells and determine whether they are targets to be eliminated. Six such molecules were quantified in the PMP replicates and showed only moderate downregulation for cadherin (CDH)2 and CDH4 (**Figure S2**). Natural cytotoxicity receptor (NCR) ligands and CD47, a ligand for signal regulatory protein alpha (SIRP-α), remained largely unchanged during VACV infection (**Figure S2**). Lastly, plexins, which are ligands for semaphorins, showed mild downregulation from the cell surface (**Figure S2**).

It is probable that not all receptor-ligand pairs involved in lymphocyte regulation have been identified. Most NK and T cell ligands display structural similarities and belong to a few protein families including cadherins, collagen, C-type lectin, TNF, MHC and immunoglobulin (40) and often these are modulated during viral infection. These characteristics were exploited to define putative candidate surface proteins with immunomodulatory functions. Host PM proteins substantially modulated during VACV infection were annotated with InterPro functional domains (41). Six upregulated and 24 downregulated human PM proteins showed InterPro domain annotations associated with NK/T cell ligands, which may influence immune recognition (**Table S3**). This included multiple protocadherins, endosialin (CD248), and several tyrosine-protein kinase receptors such as AXL, PTPRK, PTPRS and TYRO3 (**Figure 5**). Interestingly, protocadherins were also downregulated after infection with HCMV (30) and knockdown of protocadherin FAT1 in target cells led to decreased NK cell degranulation (29). Additionally, FGFR1 was reported to co-stimulate T cells (42), and targeting of AXL sensitised lung cancer cells to lymphocyte-mediated cytotoxicity (43). Taken together, these proteins modulated during VACV infection may represent putative immune ligands.

**Figure 5.**
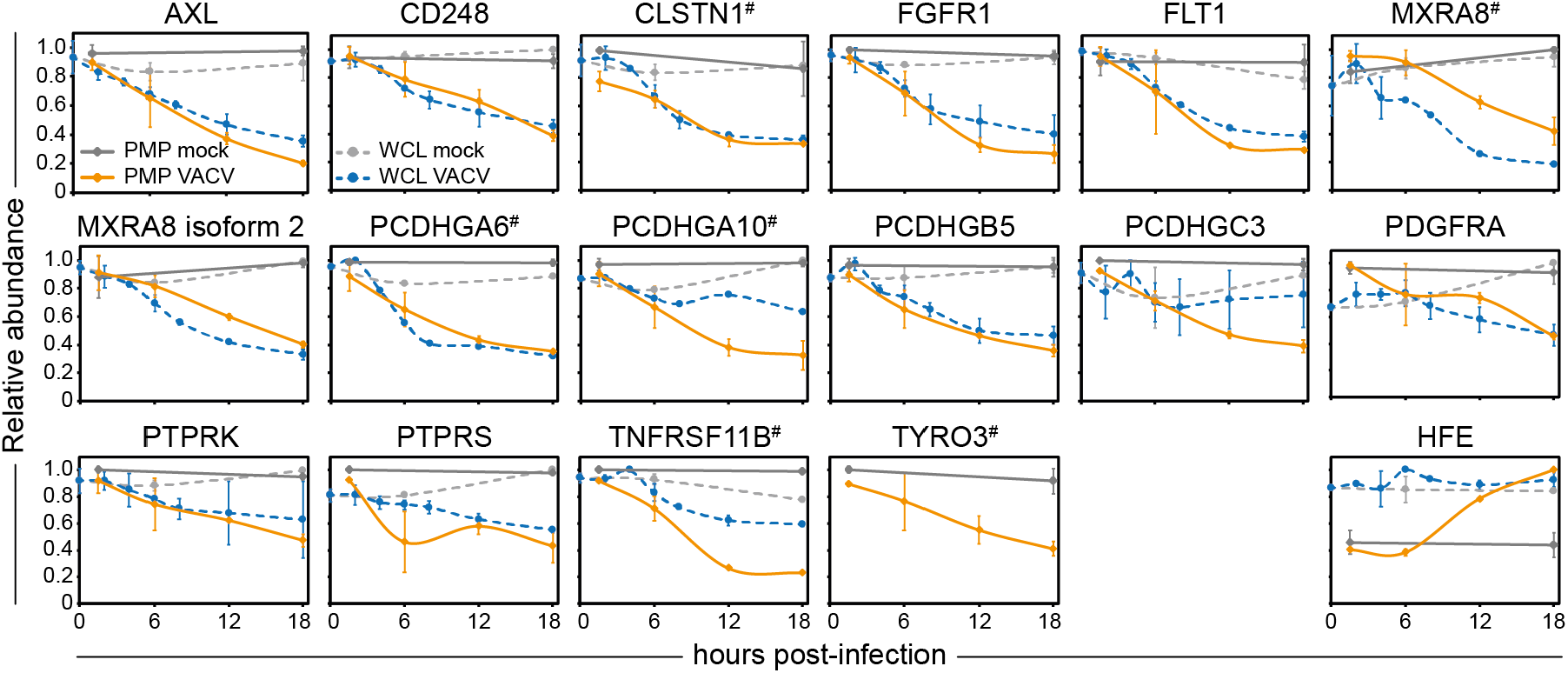
Modulation of surface expression of putative immune ligands during VACV infection. Temporal profiles of selected putative immune ligands modulated during VACV infection (**Table S3**). Data are represented as mean ± SD (PMP n=2; WCL n=3 (9), # WCL < n=3).

### VACV proteins detected at the plasma membrane

The VACV proteins detected at the PM increased in number and abundance as infection progressed (**Figure 1C-D**). Given the poor annotation for the subcellular location of many VACV proteins, a filtering strategy was applied to discriminate between VACV proteins that are likely to be true PM proteins and non-PM contaminants that may have been detected due, for example, to high intracellular abundance. The number of peptides identified for a given protein was compared between PMP and WCL (9) proteomic datasets. For each human protein quantified in both PMP replicates, a peptide count ratio was calculated (29): (peptide counts PMP 1+2) / (peptide counts whole cell lysate 1+2+3). More than 90% of the human proteins that were GO-annotated as non-PM showed a peptide ratio <0.5, whereas 85% of the proteins scoring above 0.5 were defined as human PM proteins (**Figure 6A**). This illustrates that the peptide ratio is a reliable metric to predict if a protein is likely to be expressed at the cell surface, or if it is a non-PM contaminant.

**Figure 6.**
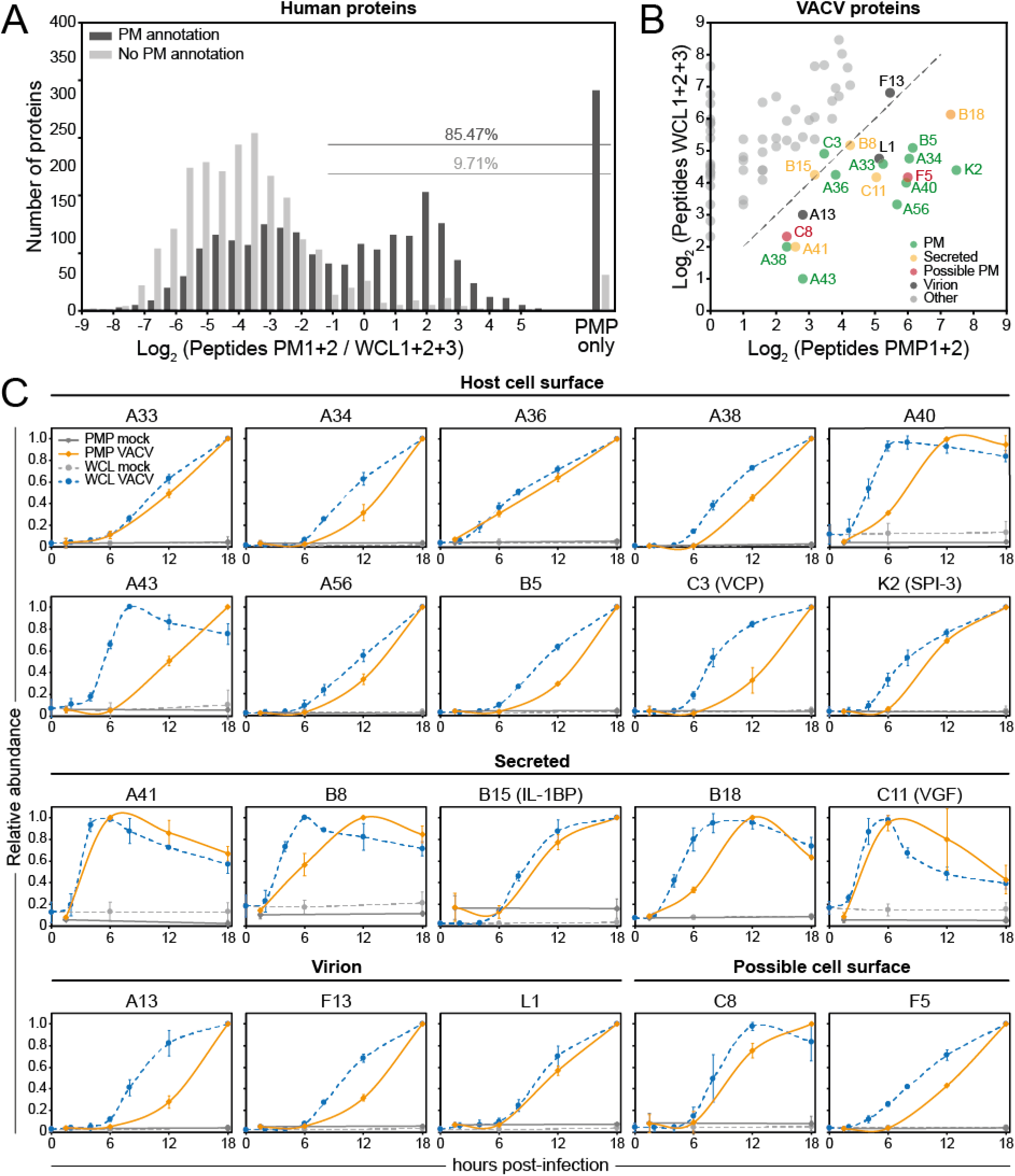
Identification of high-confidence VACV PM proteins. (**A**) Peptide ratios comparing the peptide count for a given protein in the PMP vs. WCL (9) for all human proteins quantified in both PMP replicates. ‘PMP only’ = not detected in any of the WCL replicates. ‘PM annotation’ includes GO terms PM, CS, XC and ShG. (**B**) Cut-off peptide ratio of 0.5 (dashed line), as determined in panel A, applied to 73 VACV proteins detected in either of the PMP replicates to identify high-confidence VACV PM proteins (**Table 1**). (**C**) Temporal profiles of all known and high-confidence VACV PM and secreted proteins. Data are represented as mean ± SD (PMP n=2; WCL n=3, A43: WCL n=1).

The peptide ratio of 0.5 was applied to the 73 viral proteins quantified in either PMP replicate, and this identified 20 VACV proteins as high-confidence PM proteins (**Figure 6B, Table 1**). Twelve of these are known to be present at the host cell surface (A33, A34, A36, A38, A40, A43, A56 [HA], B5, B18, C3 [VCP], C11 [VGF-1] and K2) and six are secreted proteins (A41, B8, B18, B15 [IL-1β-BP], C3 and C11) (44), illustrating the validity of the filtering strategy. Note that some of the secreted proteins are also retained at the cell surface. Five other VACV proteins were identified with high-confidence as PM proteins, including the structural proteins A13 and L1 that form part of intracellular mature virus (IMV) surface (45, 46) and F13 that is present on the internal face of the outer membrane of the extracellular enveloped virus (EEV) (47, 48). The non-structural proteins C8 and F5 have not been described to interact with other VACV proteins and were identified as putative novel VACV PM proteins. F5 was reported to be located near the cell periphery, although has not been demonstrated previously to be exposed extracellularly (49).

**Table 1.** Details of high-confidence VACV PM proteins. (related to Figure 6). Gene names from VACV strain Copenhagen (Cop) have L or R to indicate direction of transcription. IEV = intracellular enveloped virus. IMV = intracellular mature virus. EEV = extracellular enveloped virus. gp = glycoprotein. *Gene non-functional in VACV strain Copenhagen. **For a review of VACV protein function and location see (44). ***Temporal classes 1 and 2 occur before viral DNA replication.

| Uniprot | Gene name VACV-Cop | Gene name VACV-WR | Protein description / function** | Known location** | Temporal class *** (9) | Functional category (9, 51) |
| --- | --- | --- | --- | --- | --- | --- |
| P01136 | C11L | VACWR009 | C11, vaccinia growth factor, VGF | Secreted | 1 | Host interaction |
| P24770 | B8R | VACWR190 | B8, IFNγ-binding protein, gp | Secreted | 1 | Host interaction |
| P24766 | A41L | VACWR166 | A41, chemokine-binding protein, gp | secreted | 1 | Host interaction |
| P25213 | B19R | VACWR200 / B18R | B18, IFNα/β-binding protein, gp | secreted & cell surface | 2 | Host interaction |
| P24765 | A40R | VACWR165 | A40, type II membrane, gp | PM | 2 | Host interaction |
| P26671 | A43R | VACWR168 | A43, gp | PM | 2 | Host interaction |
| P18384 | K2L | VACWR033 | K2, serine protease inhibitor 3 (SPI-3) | secreted & PM | 3 | Host interaction |
| P68619 | A36R | VACWR159 | A36, protein egress and actin polymerisation | PM | 3 | Virion association (IEV only) |
| P68638 | C3L | VACWR025 | C3, vaccinia complement control protein (VCP) | secreted & cell surface | 3 | Host interaction |
| P17364 | C8L | VACWR020 | C8 | Unknown | 3 | Unknown |
| P24761 | A34R | VACWR157 | A34, EEV envelope, gp | PM | 4 | Virion association (EEV) |
| Q01227 | B5R | VACWR187 | B5, EEV envelope, gp | PM | 4 | Virion association (EEV) |
| P24358 | F5L | VACWR044 | F5, major membrane protein | Unknown | 4 | Unknown |
| Q01218 | A56R | VACWR181 | A56, haemagglutinin, EEV envelope, gp | PM | 4 | Virion association (EEV) |
| P68617 | A33R | VACWR156 | A33, EEV envelope, gp | PM | 4 | Virion association (EEV) |
| Q76ZQ4 | A13L | VACWR132 | A13, IMV membrane protein | IMV | 4 | Virion association (IMV) |
| P24763 | A38L | VACWR162 | A38, gp | PM | 4 | Host interaction |
| P25212 | B16R * | VACWR197 / B15R | B15, IL-1β-binding protein | secreted | 4 | Host interaction |
| P07612 | L1R | VACWR088 | L1, IMV membrane protein | IMV | 4 | Virion association (IMV) |
| P04021 | F13L | VACWR052 | F13, EEV membrane-associated protein | EEV | 4 | Virion association (EEV) |

VACV protein expression has been categorised into four temporal classes (9, 50-54). The high-confidence VACV PM proteins cover the four temporal classes, although the majority are expressed late during infection (temporal profile (Tp) 4, **Table 1**). Overall, the expression kinetics of the VACV proteins showed a slight delay in PM expression compared to the whole cell, which may reflect the time required for protein transport to the cell surface (**Figure 6C**). Interestingly, the temporal profiles of some secreted VACV proteins showed a reduction in abundance at late time-points, consistent with protein secretion (**Figure 6C**). Notably, the appearance of proteins C3 (VCP), and K2 (SPI-3) at the PM closely matched the kinetics of cell surface A56 (HA) to which C3 and K2 bind (20).

### Mechanisms underlying changes in human PM protein expression during VACV infection

The expression of PM proteins during viral infection can be modulated by several mechanisms such as proteasomal/lysosomal degradation, arrest of synthesis or protein translocation. MG132 inhibits proteasomal degradation but also affects lysosomal cathepsins (55). To identify which PM proteins are modulated by active degradation, a VACV-infected and a mock sample were treated with MG132. MG132 was added at 2 hpi to allow the uncoating of VACV that relies on the proteasome (56-58). An MG132 rescue ratio (RR) was calculated by comparing the abundance of a given protein during VACV infection ±MG132 with the abundance of the same protein during mock-treatment ±MG132. Of the 73 proteins downregulated >2-fold in both replicates at 18 hpi, six (8.2%) showed a RR >1.5 (**Figure 7A-B, Table S4A**). EPHB3 and APP also showed a RR >1.5 in the WCL MG132 analysis (9) (**Table S1**). NECTIN1, INFAR2 and EPHB2 were not detected in the WCL MG132 dataset. Taken together, unlike the WCL proteins, cell surface protein downregulation is likely regulated predominantly through mechanisms other than proteasomal degradation.

**Figure 7.**
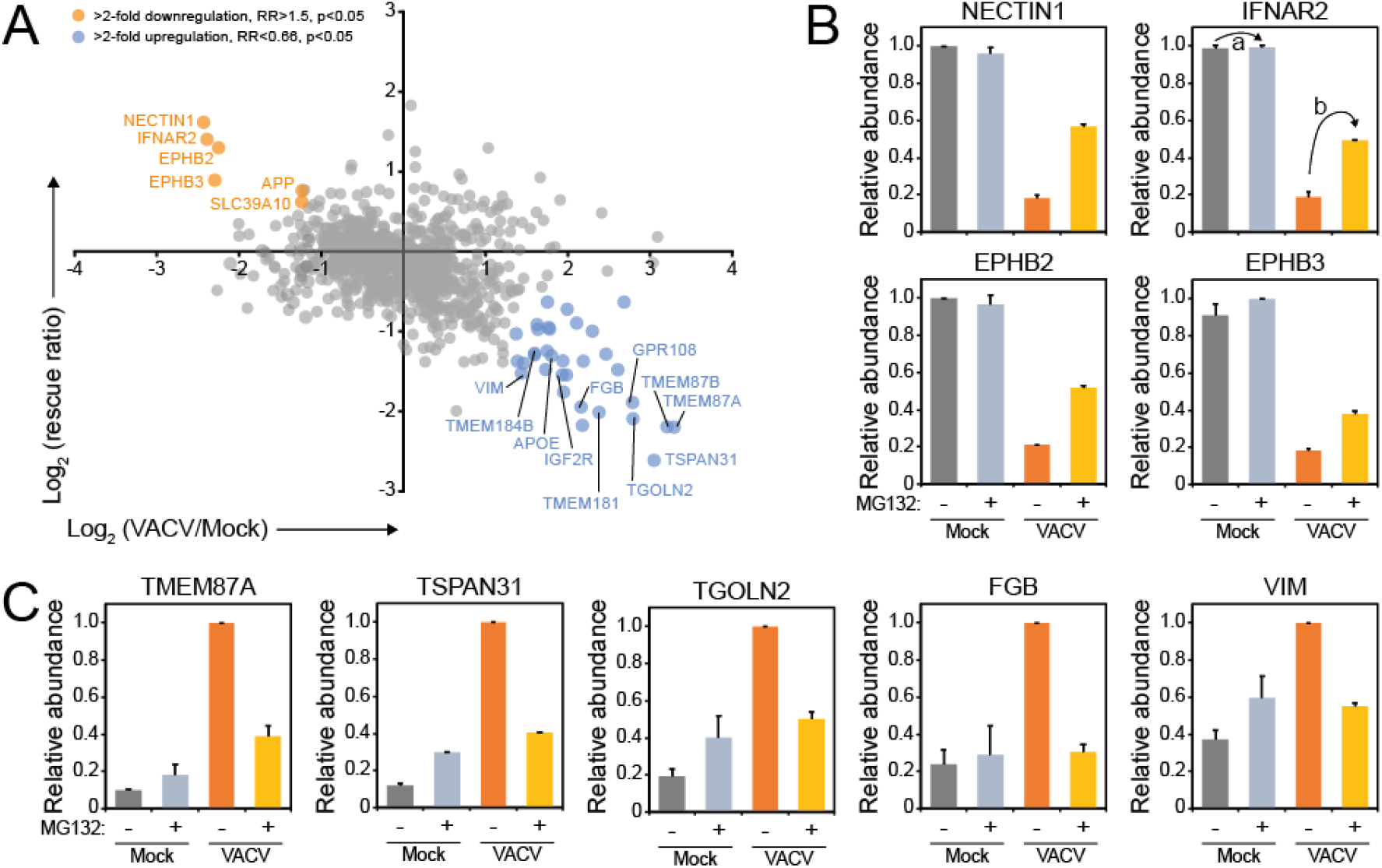
Systematic analysis of proteasome-dependent changes in surface protein expression during VACV infection. (**A**) Identification of human PM proteins downregulated from the cell surface at 18 hpi (compared to 18 h mock) in both replicates and rescued by addition of MG132 (>2 FC, rescue ratio (RR) >1.5, p<0.05), or upregulated at the cell surface at 18 hpi (compared to 18 h mock) and diminished by the addition of MG132 (>2-FC, RR<0.66, p<0.05) (**Table S4**). Here, we define the RR = b / a, where a = protein abundance during VACV infection +MG132 / abundance during infection -MG132. This value was limited to 1 to avoid artificial ratio inflation. b = protein abundance during mock-treatment +MG132 / abundance during mock -MG132 (see panel B, IFNAR2). P-values were estimated using significance A with Benjamini-Hochberg correction for multiple hypothesis testing (32). (**B**) Relative abundance of selected human proteins downregulated at the PM at 18 hpi and rescued by addition of MG132. (**C**) Relative abundance of selected human PM proteins for which upregulation was prevented by the addition of MG132 at 18hpi. Data are represented as mean ± SD (n=2). **Table S1**.

Interestingly, addition of MG132 modulated cell surface expression of approximately a third of the human PM proteins upregulated during VACV infection (**Figure 7A/C, Table S4B**). In this and previous studies, it was observed that addition of MG132 inhibits expression of late VACV genes, but not early genes (9, 57, 58). Therefore, proteasome-dependent upregulation of proteins at the cell surface may indicate that a late VACV protein is responsible for the observed increase in expression. Alternatively, these proteins may normally be retained inside the cell by a second host protein which is degraded by the proteasome during VACV infection, resulting in upregulation at the cell surface. VACV is known to shut-off host protein synthesis (59-63). Consequently, proteins with a short half-life may be downregulated from the PM during VACV infection due to natural turnover. In a previous study, protein turnover in HFFF-TERTs was quantified over 18 h by pulse (p)SILAC and 730 human PM proteins were identified (30). The abundance of 12 host proteins downregulated >2-fold from the cell surface during VACV infection was also shown to be reduced >2 fold in the pSILAC study (**Figure S3, Table S4C**). Taken together, these data suggest that proteasomal degradation and host protein synthesis shut-off are not the major mechanisms by which PM proteins are downregulated during VACV infection.

Host PM expression can be modulated during viral infection by degradation or enhanced production, but also by a translocation mechanism such as secretion, shedding, enhanced intracellular recycling or intracellular trapping. To identify human PM proteins that are up/downregulated during VACV infection via a translocation mechanism, protein expression levels in the PMP and WCL datasets were compared. The WCL dataset (9) was filtered for human proteins quantified in any of the replicates and showing on average >2 FC at any time-point (2, 4, 6, 8, 12, 18 hpi) compared to 18 h mock sample to determine which proteins are up-/downregulated during VACV infection.

Eighty-eight point five percent of the proteins downregulated >2-fold from the cell surface during VACV infection were also quantified in our prior WCL proteomic analysis, representing a total of 100 proteins. Within this group, about a third were downregulated in both studies (**Figure 8A, Table S5A-B**). DAVID functional enrichment analysis of these proteins showed significant enrichment of functional clusters, including ‘Tyrosine-protein kinase’, ‘Protein autophosphorylation’, ‘Cadherin’, ‘Heparin-binding’ and ‘Postsynaptic membrane’ (**Figure 8B, Table S5C**). This includes the ephrin protein family, IL6-ST and several, but not all, protocadherins (**Figures 2C & 8C**). An additional ∼20 human PM proteins downregulated in the PMP dataset showed a downward trend in the WCL but did not meet the cut-off criteria (e.g. EGFR or protocadherin gamma (PCDHG) A10, **Figure 2C**). The remainder of the proteins were downregulated solely at the PM, indicating internalisation without active degradation (e.g. IFNAR2 and KIT, **Figure 2C**).

**Figure 8.**
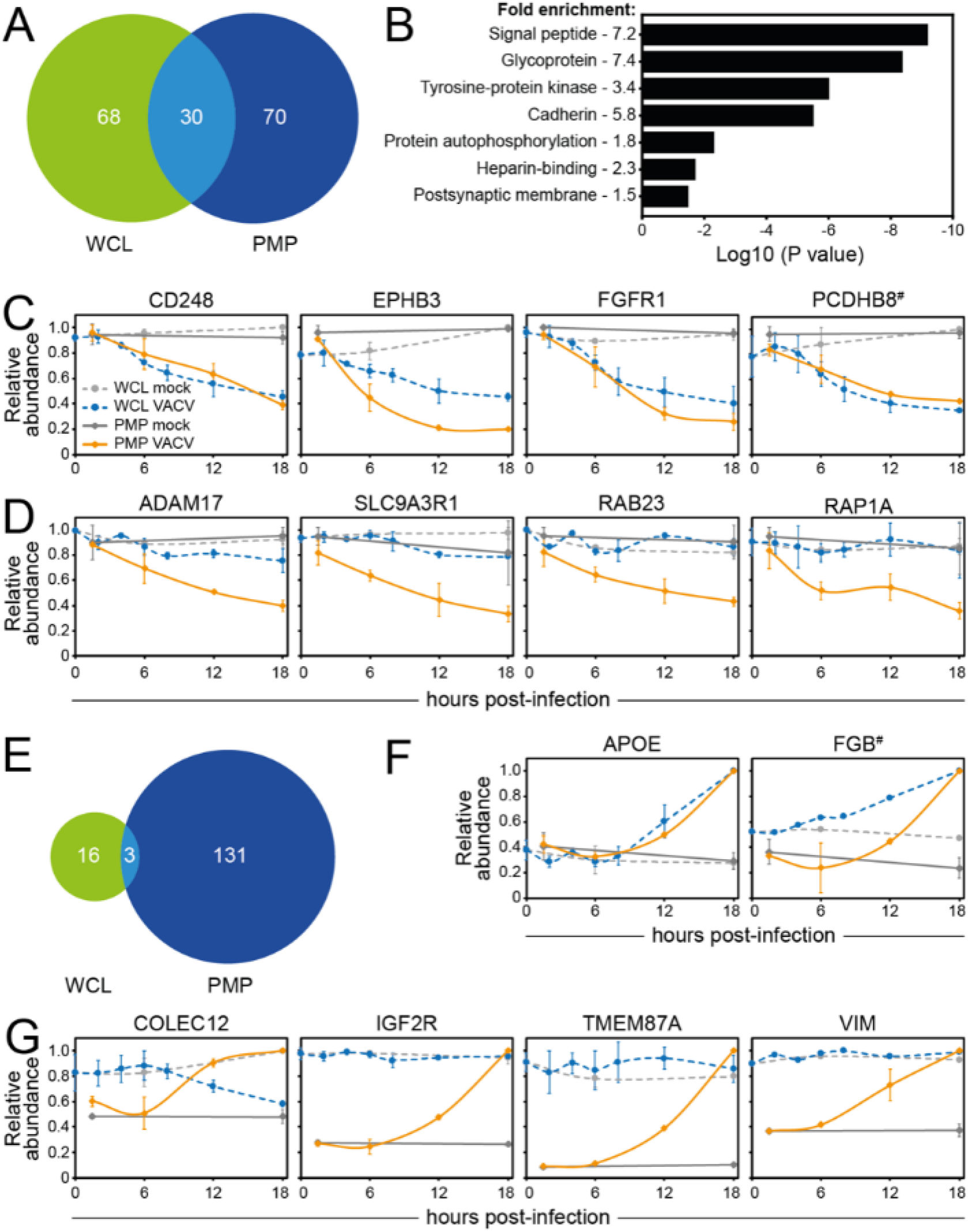
Cell surface-specific regulation of protein expression during VACV infection. (**A**) Overlap of downregulated proteins between PMP and WCL according to ‘sensitive’ criteria (**Table S5A-B**). ‘Sensitive’ criteria WCL (n=3) (9): human proteins quantified in any of the replicates and showing on average >2-fold-change at any time-point (2, 4, 6, 8, 12, 18 hpi) compared to 18 h mock sample. (**B**) DAVID functional enrichment of the 30 proteins commonly downregulated from the cell surface and at whole-cell levels. A background of all proteins detected in PMP and WCL proteomics was used. Representative terms from each cluster with a Benjamini-Hochberg-corrected p-value of <0.05 are shown (**Table S5C**). (**C**) Temporal profiles of selected host proteins that were commonly downregulated from the cell surface and at whole-cell level. (**D**) Temporal profiles of selected human PM proteins that were only downregulated at the cell surface, but not at whole-cell level (**E**) Same as panel A using proteins upregulated according to ‘sensitive’ criteria (**Table S5D**). (**F**) Temporal profiles of selected human PM proteins that were commonly upregulated from the cell surface and at whole-cell level. (**G**) Temporal profiles of selected human PM proteins that are only upregulated at the cell surface, but not at whole-cell level. Data are represented as mean ± SD (PMP n=2; WCL n=3 (9), # WCL n<3).

Ninety-four point four present of the human proteins >2-fold upregulated at the cell surface were also detected in the WCL proteomics, representing 134 proteins. Only three of these proteins were upregulated at both the cell surface and whole cell level during VACV infection: apolipoprotein E (APOE), TNFRSF10D and fibrinogen β-chain (FGB, **Figures 4B & 8E-G, Table S5D**). This is suggestive of enhanced protein synthesis, despite host shutoff. However, most upregulated proteins, including TGOLN2, HSPA5, CALR, ERAP1, SDC4, TNFRSF12A, COLEC12 and VIM, were only upregulated at the cell surface, indicating translocation to and/or stabilisation at the surface (**Figures 2E & 4B & 8F, Table S5E**).

### Identification of human PM proteins commonly targeted during virus infections

Host proteins that play an important role in antiviral immunity are often targeted by multiple viruses. To identify these proteins, the PMP dataset was compared to a published dataset analysing the cell surface proteome during infection with another dsDNA virus, HCMV (29, 64). Given the different replicative niche of VACV and HCMV, common targets of these two viruses are of particular interest. Human PM proteins quantified in either or both replicates and showing on average >2 FC at any time-point (24, 48, 72 hpi compared to the average of mock samples) were considered as up-/downregulated during HCMV infection (**Table S6**). VACV infection led to a more selective modulation of protein expression compared to HCMV, but a considerable overlap between PM proteins up- or downregulated during VACV or HCMV infection was observed (**Figure 9A/D, Table S6A/C**).

**Figure 9.**
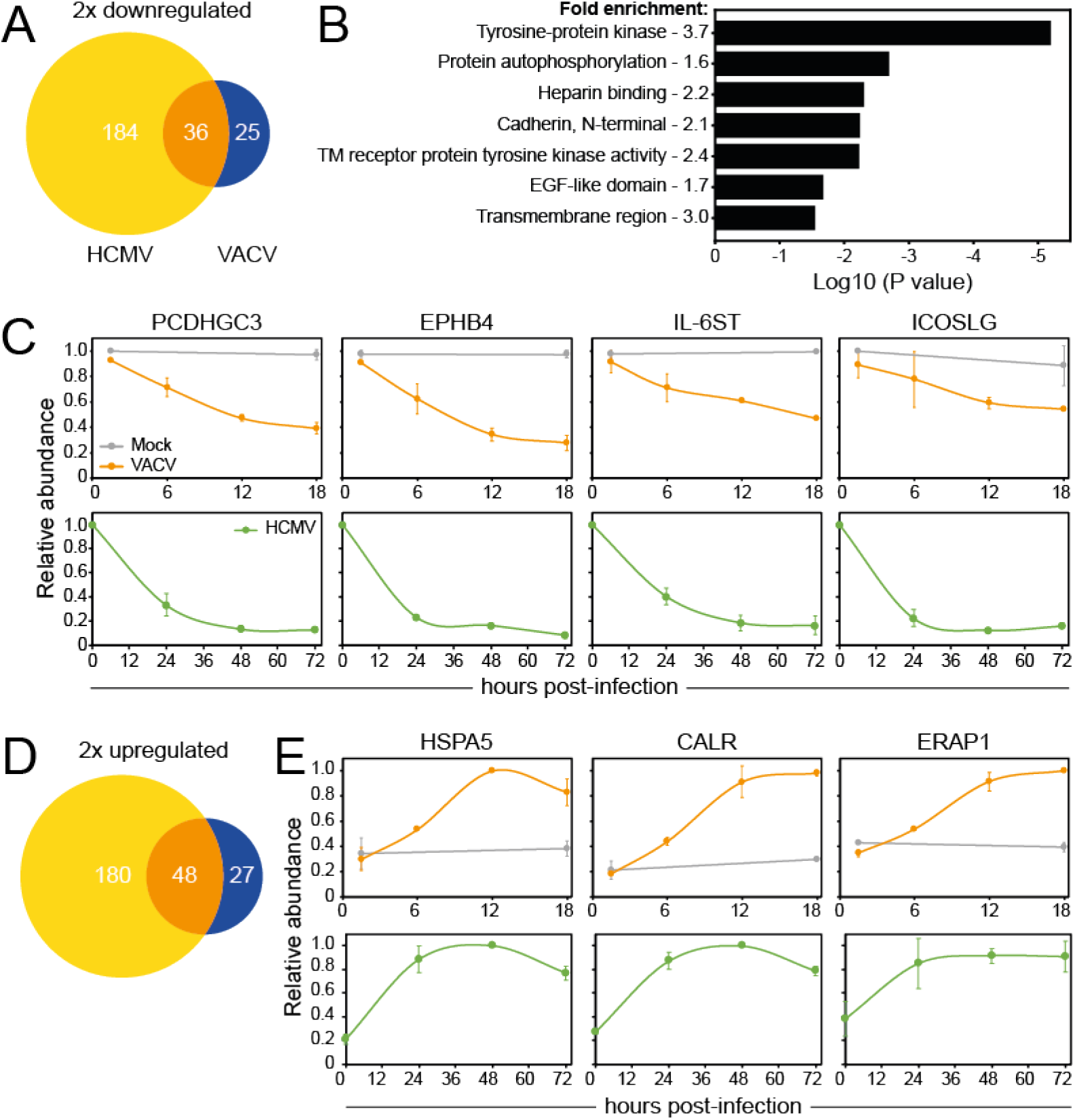
HCMV and VACV commonly target a subset of PM proteins. (**A**) Overlap of proteins downregulated according to ‘sensitive’ criteria after infection with VACV or HCMV (**Table S6A**). ‘Sensitive’ criteria HCMV PMP (n=2) (29): human PM- (GO terms PM/CS/XC/ShG) proteins quantified in a single or both replicates and showing on average >2 FC at any time-point (24, 48, 72 hpi) compared to (average of) mock sample(s). (**B**) Functional enrichment within proteins commonly downregulated from cell surface during VACV or HCMV infection (‘sensitive’ criteria). A background of all proteins detected in at least one replicate of both PMP VACV and PMP HCMV was used. Shown are representative terms from each cluster with a Benjamini-Hochberg-corrected p-value of <0.05 (**Table S6B**). (**C**) Temporal profiles of selected proteins commonly downregulated from the cell surface after VACV or HCMV infection. (**D-E**) Same as panel A/C, respectively, using proteins upregulated according to ‘sensitive’ criteria (**Table S6C**).

Fifty-four percent of the proteins downregulated at the cell surface during VACV infection were quantified in the HCMV PMP dataset. DAVID analysis of the commonly downregulated proteins revealed that both viruses target RTKs - EPHB3, EPHA4, EPHA2, EPHB4, ERBB2, PDGFRA, FLT1, ERBB4, EPHB2 -, molecules with EGF-like domains - THBS2, NID2, FBLN5, CD248, FBLN1, THBS1, VASN -, and cadherins - PCDHGB5, PCDHGA10, PCDHGC5, PCDHGC3 (**Figure 9B, Table S6B**). Other noteworthy proteins downregulated by both viruses include IL-6ST and ICOSLG, indicating that these proteins may be important in the antiviral response (**Figure 9C**).

Fifty-two point eight percent of the proteins upregulated at the PM during VACV infection were quantified in the HCMV PMP dataset. DAVID functional enrichment of proteins upregulated at the cell surface by both VACV and HCMV infection revealed enrichment of functional clusters for ‘Protein processing ER’ and ‘Prevents secretion ER’. These clusters likely relate to ER stress, which is commonly triggered during viral infections due to a substantial increase in protein production. Several of the PM proteins that were strongly upregulated during VACV infection were also upregulated at the cell surface during HCMV infection, including HSPA5, CALR and ERPA1 (**Figure 9E, Table S6C**).

## Discussion

In this study, quantitative temporal plasma membrane proteomics was used to assess systematically the impact of VACV infection on host and viral PM protein expression. Altered PM protein expression may represent a normal response to viral infection, VACV-mediated suppression of immune responses and/or modulation of the environment to support virus production and spread.

VACV infection resulted in selective modulation of host PM protein expression. Notably, substantial downregulation of several members of the RTK and protocadherin families was observed and these protein families were also downregulated from the PM during HCMV infection (29). Several members of the RTK (FGFR1, PDGFRA, KIT, FLT1, AXL, TYRO3) and protocadherin (PCDHGA10, PCDHGA6, PCDHGB5, PCDHGB7, PCDHGC3) families contain InterPro functional domains often found in immune ligands (40, 65), suggesting that they may act as immune regulators. This hypothesis is supported by previous reports that PM expression of the protocadherin FAT1 leads to decreased degranulation of NK cells (29) and that the RTK ephrin B2 leads to T cell co-stimulation (66). Particularly notable was the downregulation of the RTK EGFR, which was also observed after HSV-1 and HCMV infection (29, 67, 68). EGFR downregulation from VACV-infected cells is of particular interest because VACV also expresses a viral epidermal growth factor (called vaccinia growth factor, VGF, protein C11) that contributes to virulence (69). VGF stimulates cells surrounding the infected cell to proliferate, causing hyperplasia and mitotic bodies characteristic of orthopoxvirus pathology (21). More recently, VGF was also reported to enhance motility of infected cells to promote viral spread (22). The removal of EGFR from the surface of VACV-infected cells may represent a strategy to promote binding of VGF to EGFR on surrounding uninfected cells thereby stimulating the metabolic activity of these cells to enhance virus replication. Alternatively, the removal of EGFR from the infected cell surface may reflect internalisation of activated EGFR upon engagement with VGF, followed by degradation rather than recycling to the PM (70).

PM proteins upregulated during VACV infection included chaperone proteins whose translocation to the PM is associated with ER stress and activation of the immune system (71). Some of these proteins are manipulated by viruses to suppress immune responses. For example, CALR suppresses IFN-α production and antiviral activity in the context of hepatitis B virus infection (72), ERAP1 induces cleavage of cytokine receptors (73, 74), and syndecan-4 (SDC4) negatively regulates retinoic acid-inducible gene I (RIG-I)-mediated signalling during virus infection (75). Overall, the increased PM expression of these proteins during VACV infection may indicate previously unknown strategies by which VACV manipulates the host response to infection.

VACV is well-known to interfere with cytokine signalling by expressing soluble binding proteins, or decoy receptors, for IL-1β, TNFα, IL-18, IFN-γ and IFN-α/β (11). In addition, IFN signalling is targeted at several levels in the pathway downstream of the receptor (76, 77). The observed downregulation of IFNAR2 (type I IFN receptor) and IL10-RB (a component of multiple cytokine receptors, including type III IFNs) from the PM, may represent novel VACV strategies for evasion of the IFN response.

NK and T cells are activated by changes on the surface of the infected cell. There are contested reports that VACV infection leads to the mild downregulation of total MHC-I surface levels (14, 24-27). However, the selective downregulation of HLA-C from the cell surface observed here is consistent with a previous study relying on HLA transfection (27). HLA-A and -B represent the majority of cell surface MHC-I (78), which explains why reduction of HLA-C did not substantially affect the total MHC-I levels detected by flow cytometry. HLA subtypes have a differential impact on the immune response and whilst all HLA-C molecules are KIR ligands, HLA-A and -B mostly interact with T cell receptors. Importantly, KIRs and HLA polymorphisms are linked to infectious disease outcome (79) and if the selective modulation of HLA-C is conserved in other orthopoxviruses, such as variola virus, this might have contributed to pathogenesis of smallpox.

The activating NKG2D ligands and B7-H6 (NKp30 ligand), which are typically upregulated in response to viral infection or in cancer cells (26, 80), were not upregulated during VACV infection and this may represent a novel strategy by which VACV evades the NK cell response. In contrast, VIM, a ligand for the activating receptor NKp46, was upregulated at the PM during VACV infection, whilst the total cell level remained unchanged. VIM upregulation sensitises mycobacterium tuberculosis-infected cells to NKp46-mediated lysis (81), facilitates adenovirus type 2 transport (82) and interacts with VACV virions and facilitates their assembly (83). Taken together, this suggests that translocation of VIM to the cell surface during VACV infection may be caused by virion transport. The immune system may have evolved a strategy to detect this through NKp46-mediated recognition of VIM.

Further, several immune checkpoints were selectively modulated during VACV infection. PD-L1 was not affected during VACV infection, which is in contrast with HCMV-induced up-regulation of PD-L1 (29). Downregulation of the costimulatory molecule ICOSLG potentially represents a novel mechanism by which VACV modulates NK and T cells (84-87). Additionally, RGMB downregulation may interfere with T cell costimulatory function (88) or affect co-inhibitory pathways (89). The impact of modulation of these proteins on the immune response remains to be determined.

More VACV proteins were detected at the PM than had been described hitherto, and a filtering strategy was used to distinguish likely true viral PM proteins from those that might represent ‘overspill’ from an abundant intracellular pool. This process identified 5 possible new PM proteins: IMV envelope proteins A13 and L1 (45, 46), the EEV outer membrane protein F13 (47) and non-structural proteins C8 and F5. L1, A13 and F13 might either be expressed at the PM or alternatively, their detection there might result from interactions with other proteins. For example, the presence of F13 might reflect its interaction with B5 and A56 (90, 91). C8 and F5 are not present in IMV particles (92-94) and are not known to interact with other VACV proteins (95). F5 has a transmembrane domain and is expressed at the periphery of the infected cell, in regions in contact with neighbouring cells (49). The subcellular localisation of C8 is unknown, but it also contains a hydrophobic domain that might function as either a signal peptide or transmembrane domain. Overall, these findings justify further study of the roles of C8 and F5 during VACV infection, particularly since non-structural PM proteins may have additional roles in immune regulation.

Systematic comparison of the PMP dataset with various other datasets gave insight into mechanisms used to regulate cell surface protein expression during VACV infection. Only six downregulated PM proteins were rescued by addition of MG132 (8.2%), which contrasts with the WCL proteomics where 69% of the proteins was rescued by MG132 (9). Nevertheless, most of the proteins downregulated from the PM were also downregulated, or showed a downward trend, at the whole-cell level. A combination of VACV-induced host shut-off and high protein turnover may explain the downregulation of a few proteins. However, most downregulated human PM proteins are likely degraded through non-proteasomal mechanisms, for example, lysosomal degradation. The remainder of the proteins were downregulated specifically from the cell surface, and not at the whole-cell level, likely indicating internalisation and/or retention in intracellular compartments without active protein degradation.

Proteins from the same family were not always regulated by the same mechanism. For example, the downregulation of PM ephrin B2 and B3, but not other ephrins, was prevented by addition of MG132. Additionally, most protocadherins were downregulated in both PMP and WCL experiments, however, protocadherin-γ A10 and C3 did not show a clear downregulation at whole-cell level. Furthermore, protocadherin-γ B4 downregulation may be the result of a high protein turnover in combination with host shut-off by VACV. This may indicate that VACV has developed several mechanisms to modulate specific members of a protein family, which may have particularly important functions as novel immune ligands.

Strikingly, even though HCMV and VACV encode a similar number of genes, VACV modulates the abundance of fewer proteins compared to HCMV. Similar observations were made after comparing WCL datasets of VACV or HCMV infection (9). The greater alteration of host protein abundance by HCMV, may reflect the longer and more complex HCMV infectious cycle including the ability to enter and exit the nucleus and establish latency. Nonetheless, there is considerable overlap between proteins targeted by both viruses. Targeting of the whole protocadherin family by multiple viruses may indicate that these proteins represent previously unknown immune ligands.

The PMP technique detects changes in expression levels, rather than modifications to proteins. Therefore, immune evasion strategies involving masking, interference with receptor-ligand binding and/or molecular mimicry may not be identified using this approach. Furthermore, the use of HFFF-TERTs – allowing for direct comparison with previously published data with the same cell type (9, 29, 30, 64) – limited the detection of immune ligands that are more commonly found on professional antigen presenting cells. Nonetheless, a number of potentially novel immunomodulatory strategies by VACV were identified. Overall, the PMP dataset represents a valuable resource providing new research avenues informing on antiviral immune responses and viral immune evasion strategies, vaccine vector design and oncolytic virus therapy.

## Materials & methods

### Cells lines

Primary HFFF immortalised with human telomerase (HFFF-TERTs) (96) were grown in Dulbecco’s modified Eagle’s medium (DMEM; Gibco, Thermo Fisher Scientific, Lutterworth, UK) supplemented with 10 % foetal bovine serum (v/v; Seralab, London, UK) and 1 % penicillin/streptomycin (p/s; Gibco, Thermo Fischer, UK). BSC-1 (African green monkey cell line, ATCC CCL-26) were grown in DMEM supplemented with 10 % filtrated bovine serum (FBS; Pan Biotech UK Limited, Dorset, UK) and 1 % P/S. HeLa cells (human cervical ATCC CCL-2) and RK13 (Rabbit kidney cells, ATCC CCL37) were grown in Minimum Essential Medium (MEM; Gibco, Thermo Fischer, Lutterworth, UK) supplemented as described above for BSC-1 cells. Non-Hodgkin’s B cell line DOHH2 (kind gift from Dr Daniel Hodson) was grown in Roswell Park Memorial Institute (RPMI)-1640 medium (Gibco, Thermo Fischer, Lutterworth, UK) with 10 % FBS and 1 % p/s. All cell lines were maintained at 37°C in 5 % CO_2_. Cells were routinely checked as mycoplasma negative (MycoAlert, Lonza, UK).

### Vaccinia virus

VACV strain Western Reserve (WR) stocks were produced by infection of RK13 cells. Virus particles were released from cells by three freeze-thaw cycles and two rounds of 20 strokes of a Dounce homogenizer. Cell-free viruses were resuspended in 10 mM Tris HCl pH 9.0 and purified by sedimentation through a 36% (w/v) sucrose cushion twice (Fisher Chemical, Thermo Fisher Scientific). Purified viruses were resuspended in PBS, titrated in BSC-1 cells by plaque assay and stored at -80 °C.

### Virus infections and MG132 treatment

Immediately before infection, the culture medium on 15 cm^2^ dishes with 4 - 6.5 × 10^6^ HFFF-TERTs was replaced with infection medium (DMEM supplemented with 2% FBS and 1% p/s). Cells were mock-treated or infected at MOI 5 with VACV in 6 ml infection medium. After 90 min adsorption at 37°C, 15 ml of infection medium was added, and samples were incubated at 37°C until harvesting. For samples treated with the proteasome inhibitor MG132, 0.5 µl/ml medium (v/v; Merck, Kenilworth, U.S.A.) of 20 mM MG132 was added per flask at 2 hpi. Infections were staggered to allow for simultaneous harvesting of the indicated time-points.

### Plasma membrane profiling and TMT labelling

Plasma membrane protein labelling was performed as described (29). Briefly, cell surface sialic acid residues were oxidised and biotinylated using 1 mM sodium periodate (Thermo Scientific), 100 mM aminooxy-biotin (Biotium Inc., Fremont, U.S.A.) in dry DMSO and 10 mM aniline (Sigma-Aldrich, Merck, Dorset, U.K.) in PBS pH 6.7 (Sigma-Aldrich). After 30 min at 4 °C, the reaction was stopped by addition of glycerol (Sigma-Aldrich) at a final concentration of 1 mM. Cells were harvested into 1.6 % Triton X-100 (Fisher Scientific), 150 mM NaCl (Sigma-Aldrich), 5 mM iodoacetamide (Sigma-Aldrich) in 10 mM Tris-HCl pH 7.6 (Sigma-Aldrich) supplemented with protease inhibitor tablets (Roche, Merck). Biotinylated glycoproteins were enriched with high-affinity streptavidin agarose beads (Pierce) and washed extensively. Captured protein were denatured with SDS and urea, reduced with DTT, alkylated with iodoacetamide (IAA, Sigma) and digested on-bead with trypsin (Promega) in 200 mM HEPES (4-(2-hydroxyethyl)-1piperazineethanesulfonic acid) pH 8.5 for 3 h. The digested peptides were eluted, and each sample labelled with 56 µg of a unique TMT reagent (Thermo Fisher Scientific) in a final acetonitrile concentration of 30 % (v/v) for 1 h at room temperature. Samples were labelled as follows: TMT 126 (WT VACV 90 min), TMT 127N (WT VACV 6 h), TMT 127C (WT VACV 12 h), TMT 128N (WT VACV 18 h), TMT 128C (Mock 18 h), TMT 130N (WT VACV + MG132 18 h), TMT 130C (Mock + MG132 18 h), TMT11-131C (Mock 90 min). The reaction was quenched with hydroxylamine to a final concentration of 0.5 % (v/v). TMT labelled samples were combined at equal ratio, vacuum-centrifuged and subjected to C18 solid-phase extraction (Sep-Pak, Waters).

### HpRP Fractionation and LC-MS3

An unfractionated single-shot sample was analysed to ensure similar peptide loading across each TMT channel. The remaining TMT-labelled tryptic peptide samples were subjected to HpRP fractionation, as described (30) except that the samples were prepared as a single-set of 6 fractions. The fractions were dried and resuspended in 10 µl MS solvent (4 % MeCN/5 % formic acid) prior to LC-MS3. Data from the single-shot experiment was analysed with data from the corresponding fractions to increase the overall number of peptides quantified. Mass spectrometry data was acquired using an Orbitrap Lumos (Thermo Fisher Scientific, San Jose, CA). An Ultimate 3000 RSLC nano UHPLC equipped with a 300 µm ID x 5 mm Acclaim PepMap µ-Precolumn (Thermo Fisher Scientific) and a 75 µm ID x 50 cm 2.1 µm particle Acclaim PepMap RSLC analytical column was used. Loading solvent was 0.1% FA, analytical solvent A: 0.1% FA and B: 80% MeCN + 0.1% FA. All separations were carried out at 40°C. Samples were loaded at 5 µL/min for 5 min in loading solvent before beginning the analytical gradient. The following gradient was used: 3-7% B over 3 min, 7-37% B over 173 min, followed by a 4-min wash at 95% B and equilibration at 3% B for 15 min. Each analysis used a MultiNotch MS3-based TMT method (97, 98). The following settings were used: MS1: 380-1500 Th, 120,000 Resolution, 2 × 10^5^ automatic gain control (AGC) target, 50 ms maximum injection time. MS2: Quadrupole isolation at an isolation width of m/z 0.7, CID fragmentation (normalised collision energy (NCE) 35) with ion trap scanning in turbo mode from m/z 120, 1.5 × 10^4^ AGC target, 120 ms maximum injection time. MS3: In Synchronous Precursor Selection mode the top 10 MS2 ions were selected for HCD fragmentation (NCE 65) and scanned in the Orbitrap at 60,000 resolution with an AGC target of 1 × 10^5^ and a maximum accumulation time of 150 ms. Ions were not accumulated for all parallelisable time. The entire MS/MS/MS cycle had a target time of 3 s. Dynamic exclusion was set to +/-10 ppm for 70 s. MS2 fragmentation was trigged on precursors 5 × 10^3^ counts and above.

### Protein quantification and data processing

Mass spectra were processed as described in (9) using MassPike, a sequest-based software pipeline, through a collaborative arrangement with Professor Steven Gygi’s laboratory (Harvard Medical School). A combined database was constructed from (i) the human UniProt database (26^th^ January 2017), (ii) the VACV strain WR UniProt database (23^rd^ February 2017), (iii) common contaminants such as porcine trypsin. The combined database was concatenated with a reverse database composed of all protein sequences in reversed order. Searches were performed using a 20 ppm precursor ion tolerance, product ion tolerance was set to 0.03 Th. TMT tags on lysine residues and peptide N termini (229.162932 Da) and carbamidomethylation of cysteine residues (57.02146 Da) were set as static modifications, while oxidation of methionine residues (15.99492 Da) was set as a variable modification. To control the fraction of erroneous protein identifications, a target-decoy strategy was employed (99, 100). Peptide spectral matches (PSMs) were filtered to an initial peptide-level false discovery rate (FDR) of 1% with subsequent filtering to attain a final protein-level FDR of 1% (101, 102). PSM filtering was performed using a linear discriminant analysis, as described (103). This distinguishes correct from incorrect peptide IDs in a manner analogous to the widely used Percolator algorithm (104), though employing a distinct machine learning algorithm. The following parameters were considered: XCorr, DCn, missed cleavages, peptide length, charge state, and precursor mass accuracy. Protein assembly was guided by principles of parsimony to produce the smallest set of proteins necessary to account for all observed peptides (103). Proteins were quantified by summing TMT reporter ion counts across all matching peptide-spectral matches using ‘‘MassPike,’’ as described (97, 98). A minimum one unique or shared peptide per protein was used for quantitation.

Briefly, a 0.003 Th window around the theoretical m/z of each reporter ion (126, 127n, 127c, 128n, 128c, 129n, 129c, 130n, 130c, 131n, 131c) was scanned for ions, and the maximum intensity nearest to the theoretical m/z was used. The primary determinant of quantitation quality is the number of TMT reporter ions detected in each MS3 spectrum, which is directly proportional to the signal-to-noise (S:N) ratio observed for each ion (105). Conservatively, every individual peptide used for quantitation was required to contribute sufficient TMT reporter ions (minimum of 1375 per spectrum) so that each on its own could be expected to provide a representative picture of relative protein abundance (97). Additionally, an isolation specificity filter was employed to minimize peptide co-isolation (106). Peptide-spectral matches with poor quality MS3 spectra (more than 9 TMT channels missing and/or a combined S:N ratio of less than 275 across all TMT reporter ions) or no MS3 spectra at all were excluded from quantitation. Peptides meeting the stated criteria for reliable quantitation were then summed by parent protein, in effect weighting the contributions of individual peptides to the total protein signal based on their individual TMT reporter ion yields.

Protein quantitation values were exported for further analysis in MS Excel. For protein quantitation, reverse and contaminant proteins were removed, then each reporter ion channel was summed across all quantified proteins and normalised assuming equal protein loading across all channels. Fractional TMT signals (i.e., reporting the fraction of maximal signal observed for each protein in each TMT channel, rather than the absolute normalised signal intensity) was used for further analysis and to display in figures. This effectively corrected for differences in the numbers of peptides detected per protein. For all proteins quantified in the PMP screens, normalized S:N ratio values are presented in **Table S1** (‘Data original’ worksheet). Peptide sequences initially assigned to HLA-A, -B or -C were manually compared to reference sequences of classical HLA-I expressed by HFFF-TERTs (HLA-A11:01, -A24:02, -B35:02, -B40:02, -C02:02, and -C04:01) (107). Only the peptides matching uniquely to the reference sequence of a single subtype (-A, -B, -C) were included. The summed S:N values of these peptides was used for the relative abundance of HLA-A, HLA-B or HLA-C and are available in **Table S1** (‘Data’ worksheet).

### Validation of infection and modulation of host protein expression levels by flow cytometry

In parallel with infections for PMP, 10 cm^2^ dishes with ∼1×10^6^ cells were mock-treated or infected with VACV at MOI 5. At 15.5 hpi, cells were harvested using trypsin-EDTA (Gibco) and fixed and permeabilised using Fixation/Permeabilization Solution Kit according to the manufacturer’s instructions (BD Biosciences, San Jose, U.S.A.). Cells were stained with a mouse anti-D8 monoclonal antibody AB1.1 (108) followed by PE-conjugated goat anti-mouse IgG (BioLegend, San Diego, U.S.A.). To assess human PM protein expression levels, HFFF-TERTs or HeLa cells were mock-treated or infected with VACV at MOI 5 in DMEM 2 % FCS, 1 % P/S. Cells were harvested at 14-16 hpi cells using accutase solution as per the manufacturer’s instructions (Sigma-Aldrich). ICOSLG surface expression could not be detected by flow cytometry on HFFF-TERTs, therefore, DOHH2 cells were infected at MOI 50 with VACV in RPMI-1640 with p/s. At 1 hpi, RPMI-1640 supplemented with 2% FBS (v/v) and 1% p/s was added to resuspend cells at 500,000 cells / 100 μl. All validation of PM protein expression was performed with 2-2.5×10^5^ cells / well, stained with Zombie NIR Fixable Viability Kit (Biolegend) in combination with MHC-I (W6/32, kindly provided by Dr L. Boyle), MICA (2C10, Santa Cruz Biotechnology, Santa Cruz, CA, USA), EPHB4 (rea923, Miltenyi Biotech, Bergisch Gladbach, Germany), CD95 (DX2, Miltenyi Biotech), EGFR (528, Santa Cruz), HLA-B/C (4E, kindly provided by Dr L. Boyle), HLA-C/E (DT9, kindly provided by Dr L. Boyle), ULBP3 (166510, R&D systems), B7-H6 (875001, R&D systems, Minneapolis, U.S.A), ULBP-2/5/6 (65903, R&D systems), Plexin-B1 (rea728, Miltenyi Biotech), MICB (236511, R&D systems), trail-R4 (104918, R&D systems), APC-conjugated ICOSLG Monoclonal Antibody (MIH12), isotype mouse Igg2b (Santa Cruz), isotype mouse IgG2a (Santa Cruz) isotype mouse IgG1 (Santa Cruz), Mouse IgG1 kappa Isotype Control (P3.6.2.8.1) conjugated with APC (eBioscience – part of Thermo Fisher Scientific, San Diego, CA, USA). Samples stained with unconjugated primary antibodies were subsequently probed with PE-conjugated Goat anti-mouse IgG (BioLegend). Cells were fixed using Fixation/Permeabilization Solution Kit according to the manufacturer’s instructions (BD Biosciences). All samples were analysed by flow cytometry using the Invitrogen Attune NxT and FlowJo software.

### Data analysis and bioinformatics

Analyses were performed using original python code using NumPy (109), pandas (110), Matplotlib (111), SciPy.stats (112) and sklearn.metrics (113) packages. To describe human PM proteins in this study, host proteins were filtered for relevant gene ontology (GO) annotations indicative of PM location; PM, ‘cell surface’ [CS], ‘extracellular’ [XC] and ‘short GO’ [ShG, 4-part term containing ‘integral to membrane’, ‘intrinsic to membrane’, ‘membrane part’, ‘cell part’ or a 5-part term additionally containing ‘membrane’]. Two sets of criteria were defined to determine which human PM proteins showed modulated expression levels during VACV infection. First, ‘sensitive’ criteria included proteins quantified in either or both PMP replicates showing >2x FC at any time-point during infection. Second, ‘stringent’ criteria limited the false discovery rate further and included proteins detected in both PMP replicates showing >2x FC with a p-value <0.05 (Benjamini-Hochberg corrected one-way ANOVA). To include all biologically relevant proteins, ‘sensitive’ criteria were used for subsequent analyses. A hierarchical clustering analysis based on uncentered Pearson correlation was performed on the FC with Cluster 3.0 (Stanford University). To avoid skewing the data with outliers, the FC was limited to 50 in either direction. A heatmap was visualised using Java TreeView (**Figure 1C**). Volcano plots in **Figure 1D** do not show VACV protein H7 because there was no quantitative value available in the mock sample. Scale of the x-axis of these plots was limited from -7 to 7, which excluded VACV proteins C6 and RPO7 from this graph (**Figure 1D & S1D**). These two proteins were considered contaminants based on their function and subcellular localisation. Pathway enrichment analyses were performed using DAVID version 6.8 (33, 34). Indicated proteins were searched against a background of all human proteins quantified using default settings. InterPro domain annotations (65) were added to proteins modulated according to ‘sensitive’ criteria (**Table S3**). Putative immune ligands were defined by domain annotations cadherin, collagen, MHC, C-type lectin, immunoglobulin, Ig, TNF or butyrophylin.

### Statistical analysis

Figures 1D, 7A, S1D: p-values were estimated using significance A with Benjamini-Hochberg correction for multiple hypothesis testing (32). For proteins quantified in both replicates, a one-way ANOVA was used to estimate p-values for FC during the infection time-course. Per replicate, the average value of mock 1.5 & 18 hpi was used as a control. P-values were corrected for multiple hypothesis testing using the Benjamini-Hochberg method. A corrected p-value of <0.05 was considered statistically significant.

### Data availability

The mass spectrometry proteomics data will be deposited to the ProteomeXchange Consortium (http://www.proteomexchange.org/) via the PRIDE partner repository. Data for all quantified VACV and human proteins are available in **Table S1** in which the ‘Plotter’ worksheet allows for interactive generation of temporal profiles.

## Acknowledgements

We would like to thank the Flow Cytometry core facility of the School of Biological Sciences, University of Cambridge for their technical assistance. The authors thank Dr Louise Boyle for HLA antibodies and Dr Daniel Hodson for DOHH2 cells. The authors acknowledge Henrietta Lacks for the HeLa cells. This work was supported by grant 090315 from the Wellcome Trust to GLS, and MR/M019810/1 from URKI MRC, and a Wellcome Senior Clinical Research Fellowship (108070/Z/15/Z) to MPW.

## Supplementary information

**Figure S1.**
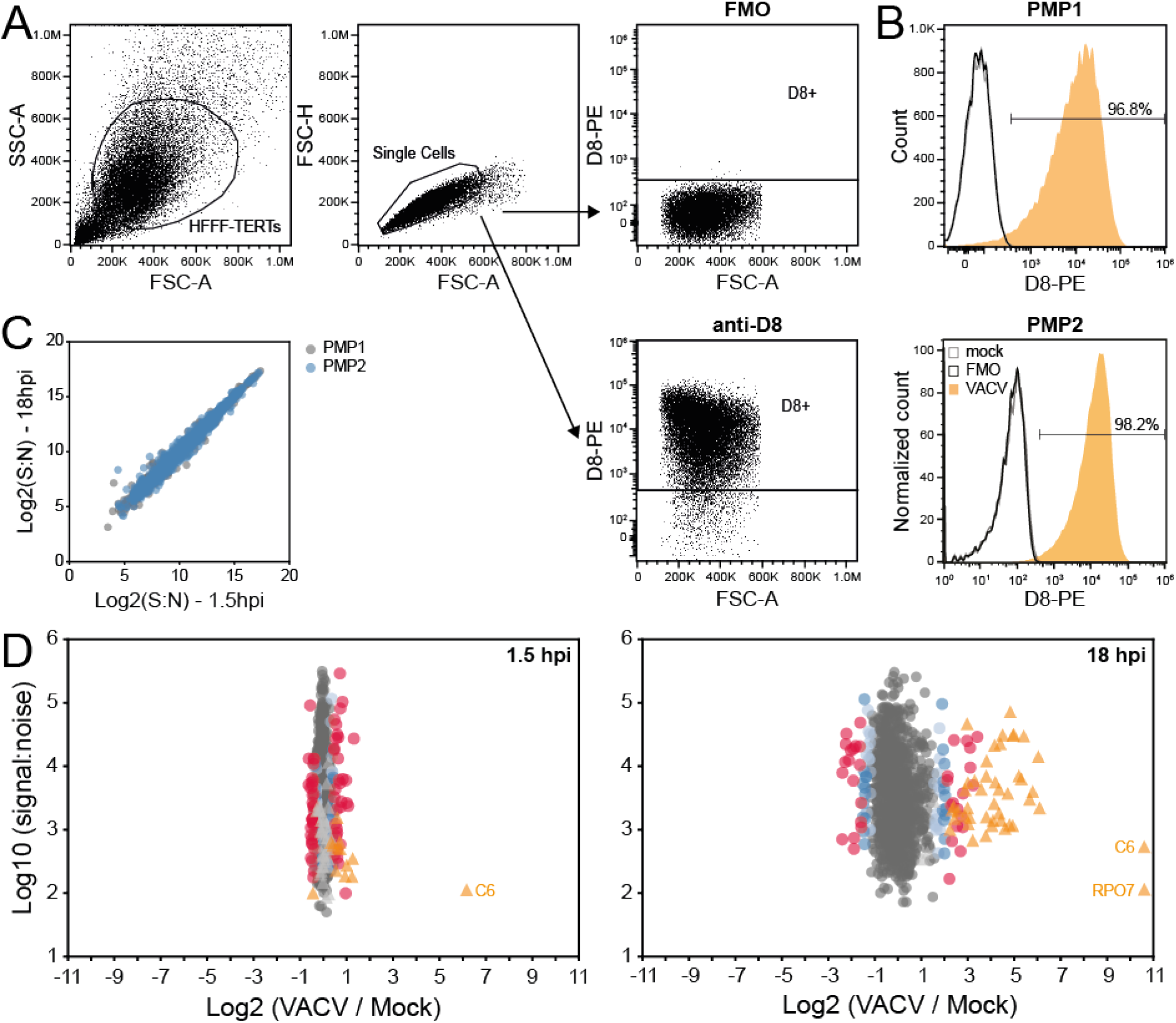
Technical details of the proteomic plasma membrane profiling experiments. Related to Figure 1. (**A-B**) HFFF-TERTs were mock-treated or infected with VACV at MOI 5 in parallel with infections for PMP. At 15.5 hpi samples were fixed and stained for the late VACV protein D8 to assess infection levels. (**A**) Representative gating strategy of VACV-infected cells stained with anti-D8 followed by anti-mouse-PE or with the secondary antibody only (fluorescence minus one, FMO) as a control. Viable cells and single cells were gated followed by selection of D8-positive cells. (**B**) D8 levels in mock-treated or VACV-infected cells for each of the two biological repeats. (**C**) Correlation of protein abundance (signal: noise, S: N) of mock-treated samples at 1.5 h and 18 h per replicate. A single human protein was excluded from PMP2 because the abundance in mock samples was ‘0’. (**D**) Fold-change of VACV and human PM proteins quantified in both repeats. Scale of the x-axis was not limited (as in **Figure 1D**) to include VACV proteins C6 and RPO7, which are considered outliers based on their function and subcellular localisation.

**Figure S2.**
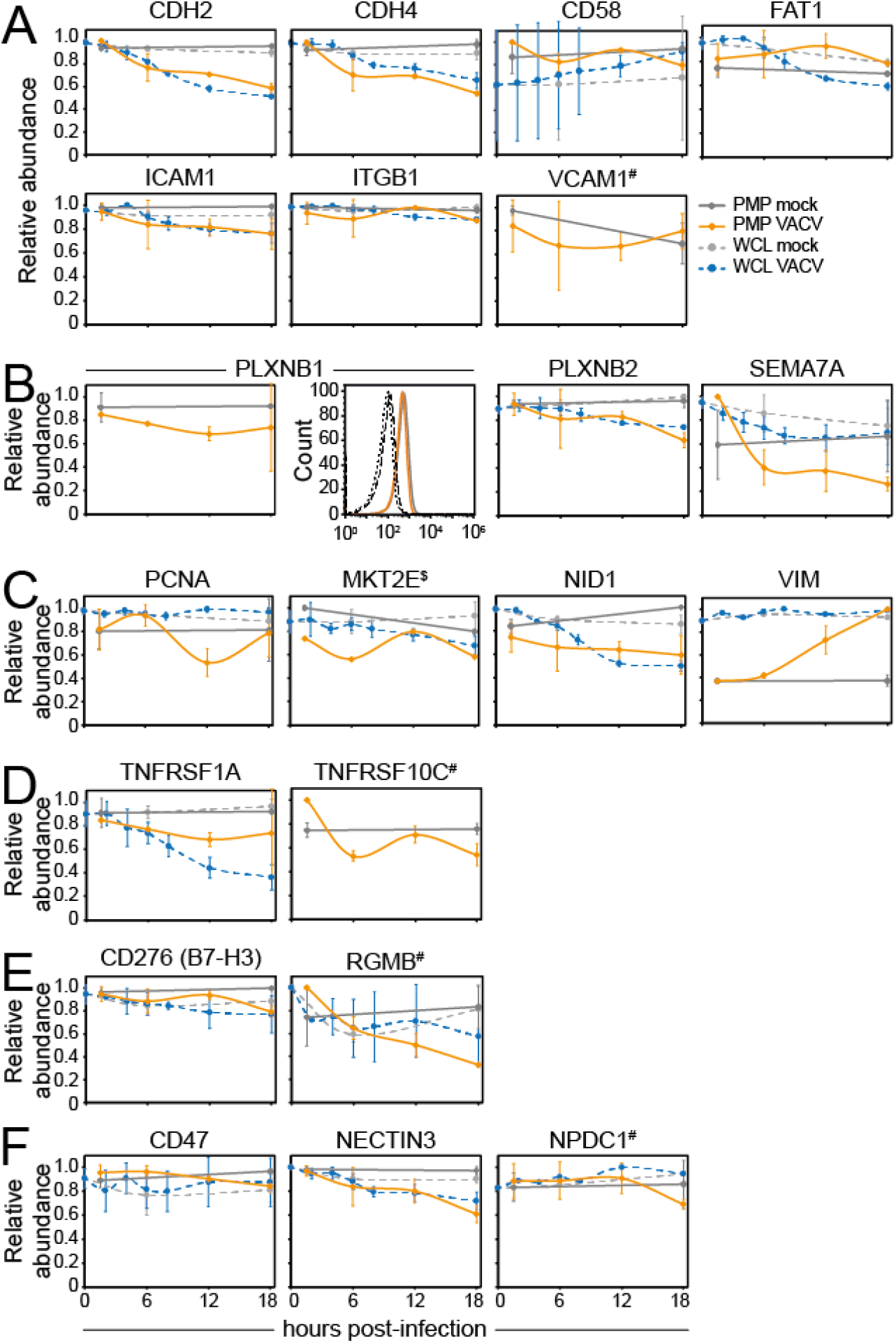
Cell surface expression of NK/T cell ligands during VACV infection. Related to Figure 4. Temporal profiles of known NK/T cell ligands. (**A**) Adhesion molecules. (**B**) Plexins. (**C**) Natural cytotoxicity triggering receptor (NCR) ligands. (**D**) Apoptosis regulators. (**E**) Co-inhibitory/stimulatory molecules. (**F**) Other. Data are represented as mean ± SD (PMP n=2, $ PMP n=1; WCL (9) n=3, # WCL < n=3). (**B**) Downregulation of plexin B1 during VACV infection was confirmed by flow cytometry in HeLa cells at 15 hpi with VACV (MOI 5). Results are representative of at least 2 independent experiments.

**Figure S3.**
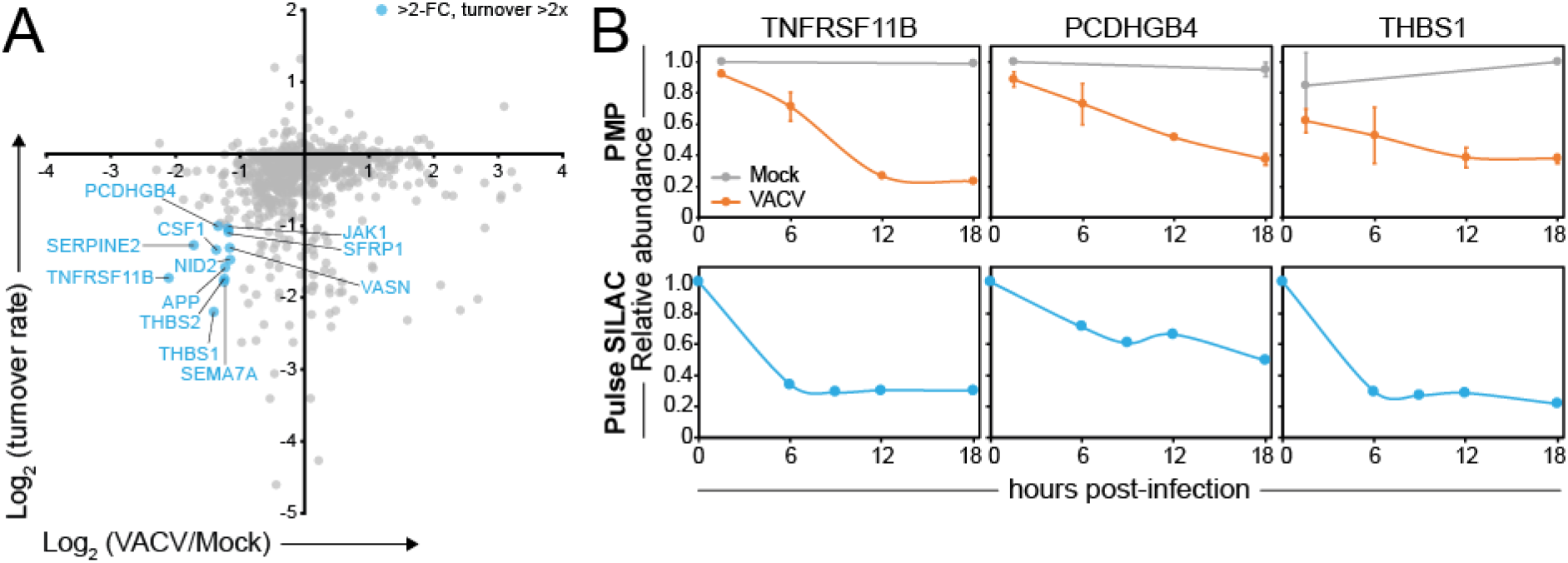
Downregulation of human PM proteins due to high protein turnover. (**A**) Identification of human PM proteins >2-fold downregulated at 18 hpi after VACV infection (compared to 18 h mock) and for which the protein abundance decreased >2-fold in 18 h was determined in a previous pulse (p)SILAC screen in mock-treated HFFF-TERTs (30). (**B**) Temporal profile of selected host proteins downregulated from the cell surface during VACV infection and which have a short half-life. Data are represented as mean ± SD (PMP n=2; pSILAC n=1 (30)).

**Table S1. Interactive spreadsheet of all data in the manuscript**. Related to Figures 2-9 & S2. The ‘Plotter’ worksheet enables the generation of graphs for all human and viral proteins quantified. The ‘Data original’ worksheet contains minimally annotated protein data where the raw data has been modified by formatting and normalisation. The ‘Data’ worksheet the HLA-A, -B, -C data is based only on peptides uniquely attributed to these three HLA subtypes. The spreadsheets include data obtained in previously performed WCL proteomics (9).

**Table S2. Plasma membrane proteins modulated during VACV infection**. Related to Figure 2. (**A-D**) Human PM proteins (A/C) downregulated or (B/D) upregulated according to (A-B) ‘sensitive’ or (C-D) ‘stringent’ criteria. (**E-F**) DAVID functional enrichment analysis of proteins shown in (A) or (B), respectively, compared to all quantified human PM proteins.

**Table S3. Discovery of putative immune ligands using InterPro domain annotation**. Related to Figure 5. InterPro functional domain annotation of proteins (A) downregulated or (B) upregulated according to ‘sensitive’ criteria.

**Table S4. Systematic analysis of mechanism underlying protein modulation during VACV infection**. Related to Figures 7&S3. (**A**) Human PM proteins quantified in both repeats and on average >2-fold downregulated compared to 18 h mock, in combination with p-value <0.05 and RR >1.5. (**B**) Human PM proteins quantified in both repeats and on average >2-fold upregulated compared to 18 h mock, in combination with p-value <0.05 and RR <0.66. (**C**) Human PM proteins quantified in both repeats and on average >2-fold downregulated compared to 18 h mock, in combination with p-value <0.05 and >2-fold pSILAC turnover rate in 18 h (30). P-values on the fold-change at 18 hpi was estimated using the method of significance A and corrected for multiple hypothesis testing (32).

**Table S5. Comparison of protein expression at whole cell level and at the plasma membrane during VACV infection**. Related to Figure 8. (**A**) Human PM proteins downregulated in both PMP and WCL experiments (9) using ‘sensitive’ criteria. (**B**) Human PM proteins downregulated in PMP but not WCL using ‘sensitive’ criteria. (**C**) DAVID functional enrichment analysis of proteins shown in (A), compared to all human PM proteins quantified in at least one replicate of PMP and WCL. (**D**) Human PM proteins upregulated in both PMP and WCL using ‘sensitive’ criteria. (**E**) Human PM proteins upregulated in PMP but not WCL using ‘sensitive’ criteria.

**Table S6. Human PM protein regulation by VACV and HCMV**. Related to Figure 9. (**A/C**) All human PM proteins (A) downregulated or (C) upregulated >2-fold by both VACV and HCMV (29) according to ‘sensitive’ criteria. (**B/D**) DAVID functional enrichment of proteins shown in (A) and (C), respectively, compared to all proteins quantified in both VACV and HCMV PMP screens.

## References

1. Fenner F, Henderson DA, Arita I, Jezek Z, Ladnyi ID. Smallpox and its eradication. World Health Organisation, Geneva. 1988.

2. Mackett M, Smith GL, Moss B. Vaccinia virus: a selectable eukaryotic cloning and expression vector. Proc Natl Acad Sci U S A. 1982;79(23):7415–9.

3. Panicali D, Paoletti E. Construction of poxviruses as cloning vectors: insertion of the thymidine kinase gene from herpes simplex virus into the DNA of infectious vaccinia virus. Proc Natl Acad Sci U S A. 1982;79(16):4927–31.

4. Moss B. Genetically engineered poxviruses for recombinant gene expression, vaccination, and safety. Proc Natl Acad Sci U S A. 1996;93(21):11341–8.

5. Altenburg AF, Kreijtz JH, de Vries RD, Song F, Fux R, Rimmelzwaan GF, et al. Modified vaccinia virus ankara (MVA) as production platform for vaccines against influenza and other viral respiratory diseases. Viruses. 2014;6(7):2735–61.

6. Prow NA, Jimenez Martinez R, Hayball JD, Howley PM, Suhrbier A. Poxvirus-based vector systems and the potential for multi-valent and multi-pathogen vaccines. Expert Rev Vaccines. 2018;17(10):925–34.

7. Lundstrom K. New frontiers in oncolytic viruses: optimizing and selecting for virus strains with improved efficacy. Biologics. 2018;12:43–60.

8. Torres-Dominguez LE, McFadden G. Poxvirus oncolytic virotherapy. Expert Opin Biol Ther. 2019;19(6):561–73.

9. Soday L, Lu Y, Albarnaz JD, Davies CTR, Antrobus R, Smith GL, et al. Quantitative temporal proteomic analysis of vaccinia virus infection reveals regulation of histone deacetylases by an interferon antagonist. Cell Rep. 2019;27(6):1920–33 e7.

10. Lu Y, Stuart JH, Talbot-Cooper C, Agrawal-Singh S, Huntly B, Smid AI, et al. Histone deacetylase 4 promotes type I interferon signaling, restricts DNA viruses, and is degraded via vaccinia virus protein C6. Proc Natl Acad Sci U S A. 2019;116(24):11997–2006.

11. Smith GL, Benfield CTO, Maluquer de Motes C, Mazzon M, Ember SWJ, Ferguson BJ, et al. Vaccinia virus immune evasion: mechanisms, virulence and immunogenicity. J Gen Virol. 2013;94(Pt 11):2367–92.

12. Veyer DL, Carrara G, Maluquer de Motes C, Smith GL. Vaccinia virus evasion of regulated cell death. Immunol Lett. 2017;186:68–80.

13. Albarnaz JD, Torres AA, Smith GL. Modulating vaccinia virus immunomodulators to improve immunological memory. Viruses. 2018;10(3).

14. Jarahian M, Fiedler M, Cohnen A, Djandji D, Hammerling GJ, Gati C, et al. Modulation of NKp30- and NKp46-mediated natural killer cell responses by poxviral hemagglutinin. PLoS Pathog. 2011;7(8):e1002195.

15. Wilcock D, Duncan SA, Traktman P, Zhang WH, Smith GL. The vaccinia virus A4OR gene product is a nonstructural, type II membrane glycoprotein that is expressed at the cell surface. J Gen Virol. 1999;80:2137–48.

16. Alcami A, Symons JA, Smith GL. The vaccinia virus soluble alpha/beta interferon (IFN) receptor binds to the cell surface and protects cells from the antiviral effects of IFN. J Virol. 2000;74(23):11230–9.

17. Montanuy I, Alejo A, Alcami A. Glycosaminoglycans mediate retention of the poxvirus type I interferon binding protein at the cell surface to locally block interferon antiviral responses. FASEB J. 2011;25(6):1960–71.

18. Kleinpeter P, Remy-Ziller C, Winter E, Gantzer M, Nourtier V, Kempf J, et al. By binding CD80 and CD86, the vaccinia virus M2 protein blocks their interactions with both CD28 and CTLA4 and potentiates CD80 binding to PD-L1. J Virol. 2019;93(11).

19. Wang X, Piersma SJ, Elliott JI, Errico JM, Gainey MD, Yang L, et al. Cowpox virus encodes a protein that binds B7.1 and B7.2 and subverts T cell costimulation. Proc Natl Acad Sci U S A. 2019;116(42):21113–9.

20. DeHaven BC, Gupta K, Isaacs SN. The vaccinia virus A56 protein: a multifunctional transmembrane glycoprotein that anchors two secreted viral proteins. J Gen Virol. 2011;92(Pt 9):1971–80.

21. Buller RML, Chakrabarti S, Moss B, Frederickson T. Cell proliferative response to vaccinia virus is mediated by VGF. Virology. 1988;164:182–92.

22. Beerli C, Yakimovich A, Kilcher S, Reynoso GV, Flaschner G, Muller DJ, et al. Vaccinia virus hijacks EGFR signalling to enhance virus spread through rapid and directed infected cell motility. Nat Microbiol. 2019;4(2):216–25.

23. Smith GL, Vanderplasschen A, Law M. The formation and function of extracellular enveloped vaccinia virus. J Gen Virol. 2002;83(Pt 12):2915–31.

24. Baraz L, Khazanov E, Condiotti R, Kotler M, Nagler A. Natural killer (NK) cells prevent virus production in cell culture. Bone Marrow Transplant. 1999;24(2):179–89.

25. Brooks CR, Elliott T, Parham P, Khakoo SI. The inhibitory receptor NKG2A determines lysis of vaccinia virus-infected autologous targets by NK cells. J Immunol. 2006;176(2):1141–7.

26. Chisholm SE, Reyburn HT. Recognition of vaccinia virus-infected cells by human natural killer cells depends on natural cytotoxicity receptors. J Virol. 2006;80(5):2225–33.

27. Kirwan S, Merriam D, Barsby N, McKinnon A, Burshtyn DN. Vaccinia virus modulation of natural killer cell function by direct infection. Virology. 2006;347(1):75–87.

28. Weekes MP, Antrobus R, Talbot S, Hor S, Simecek N, Smith DL, et al. Proteomic plasma membrane profiling reveals an essential role for gp96 in the cell surface expression of LDLR family members, including the LDL receptor and LRP6. J Proteome Res. 2012;11(3):1475–84.

29. Weekes MP, Tomasec P, Huttlin EL, Fielding CA, Nusinow D, Stanton RJ, et al. Quantitative temporal viromics: an approach to investigate host-pathogen interaction. Cell. 2014;157(6):1460–72.

30. Nightingale K, Lin KM, Ravenhill BJ, Davies C, Nobre L, Fielding CA, et al. High-definition analysis of host protein stability during human cytomegalovirus infection reveals antiviral factors and viral evasion mechanisms. Cell Host Microbe. 2018;24(3):447–60 e11.

31. Hsu JL, van den Boomen DJ, Tomasec P, Weekes MP, Antrobus R, Stanton RJ, et al. Plasma membrane profiling defines an expanded class of cell surface proteins selectively targeted for degradation by HCMV US2 in cooperation with UL141. PLoS Pathog. 2015;11(4):e1004811.

32. Cox J, Mann M. MaxQuant enables high peptide identification rates, individualized p.p.b.-range mass accuracies and proteome-wide protein quantification. Nat Biotechnol. 2008;26(12):1367–72.

33. Huang da W, Sherman BT, Lempicki RA. Systematic and integrative analysis of large gene lists using DAVID bioinformatics resources. Nat Protoc. 2009;4(1):44–57.

34. Huang da W, Sherman BT, Lempicki RA. Bioinformatics enrichment tools: paths toward the comprehensive functional analysis of large gene lists. Nucleic Acids Res. 2009;37(1):1–13.

35. Linger RM, Keating AK, Earp HS, Graham DK. TAM receptor tyrosine kinases: biologic functions, signaling, and potential therapeutic targeting in human cancer. Adv Cancer Res. 2008;100:35–83.

36. Parham P, Norman PJ, Abi-Rached L, Guethlein LA. Human-specific evolution of killer cell immunoglobulin-like receptor recognition of major histocompatibility complex class I molecules. Philos Trans R Soc Lond B Biol Sci. 2012;367(1590):800–11.

37. Brandt CS, Baratin M, Yi EC, Kennedy J, Gao Z, Fox B, et al. The B7 family member B7-H6 is a tumor cell ligand for the activating natural killer cell receptor NKp30 in humans. J Exp Med. 2009;206(7):1495–503.

38. Matta J, Baratin M, Chiche L, Forel JM, Cognet C, Thomas G, et al. Induction of B7-H6, a ligand for the natural killer cell-activating receptor NKp30, in inflammatory conditions. Blood. 2013;122(3):394–404.

39. Raulet DH, Gasser S, Gowen BG, Deng W, Jung H. Regulation of ligands for the NKG2D activating receptor. Annu Rev Immunol. 2013;31:413–41.

40. Vivier E, Tomasello E, Baratin M, Walzer T, Ugolini S. Functions of natural killer cells. Nat Immunol. 2008;9(5):503–10.

41. Blum M, Chang HY, Chuguransky S, Grego T, Kandasaamy S, Mitchell A, et al. The InterPro protein families and domains database: 20 years on. Nucleic Acids Res. 2021;49(D1):D344–D54.

42. Kos FJ, Chin CS. Costimulation of T cell receptor-triggered IL-2 production by Jurkat T cells via fibroblast growth factor receptor 1 upon its engagement by CD56. Immunol Cell Biol. 2002;80(4):364–9.

43. Terry S, Abdou A, Engelsen AST, Buart S, Dessen P, Corgnac S, et al. AXL targeting overcomes human lung cancer cell resistance to NK- and CTL-mediated cytotoxicity. Cancer Immunol Res. 2019;7(11):1789–802.

44. Moss B, Smith GL. Poxviridae: the viruses and their replication. In: Howley PM, Knipe DM, editors. Fields Virology: DNA viruses. 2. 7th ed: Wolters Kluwer Inc; 2021. p. 573–613.

45. Jensen ON, Houthaeve T, Shevchenko A, Cudmore S, Ashford T, Mann M, et al. Identification of the major membrane and core proteins of vaccinia virus by two-dimensional electrophoresis. J Virol. 1996;70(11):7485–97.

46. Martin KH, Grosenbach DW, Franke CA, Hruby DE. Identification and analysis of three myristylated vaccinia virus late proteins. J Virol. 1997;71(7):5218–26.

47. Hirt P, Hiller G, Wittek R. Localization and fine structure of a vaccinia virus gene encoding an envelope antigen. J Virol. 1986;58(3):757–64.

48. Blasco R, Moss B. Extracellular vaccinia virus formation and cell-to-cell virus transmission are prevented by deletion of the gene encoding the 37,000 Dalton outer envelope protein. J Virol. 1991;65:5910–20.

49. Dobson BM, Procter DJ, Hollett NA, Flesch IE, Newsome TP, Tscharke DC. Vaccinia virus F5 is required for normal plaque morphology in multiple cell lines but not replication in culture or virulence in mice. Virology. 2014;456-457:145–56.

50. Yang Z, Reynolds SE, Martens CA, Bruno DP, Porcella SF, Moss B. Expression profiling of the intermediate and late stages of poxvirus replication. J Virol. 2011;85(19):9899–908.

51. Yang Z, Bruno DP, Martens CA, Porcella SF, Moss B. Simultaneous high-resolution analysis of vaccinia virus and host cell transcriptomes by deep RNA sequencing. Proc Natl Acad Sci U S A. 2010;107(25):11513–8.

52. Assarsson E, Greenbaum JA, Sundstrom M, Schaffer L, Hammond JA, Pasquetto V, et al. Kinetic analysis of a complete poxvirus transcriptome reveals an immediate-early class of genes. Proc Natl Acad Sci U S A. 2008;105(6):2140–5.

53. Croft NP, de Verteuil DA, Smith SA, Wong YC, Schittenhelm RB, Tscharke DC, et al. Simultaneous quantification of viral antigen expression kinetics using data-independent (DIA) mass spectrometry. Mol Cell Proteomics. 2015;14(5):1361–72.

54. Yang Z, Cao S, Martens CA, Porcella SF, Xie Z, Ma M, et al. Deciphering poxvirus gene expression by RNA sequencing and ribosome profiling. J Virol. 2015;89(13):6874–86.

55. Wiertz EJ, Jones TR, Sun L, Bogyo M, Geuze HJ, Ploegh HL. The human cytomegalovirus US11 gene product dislocates MHC class I heavy chains from the endoplasmic reticulum to the cytosol. Cell. 1996;84(5):769–79.

56. Mercer J, Snijder B, Sacher R, Burkard C, Bleck CK, Stahlberg H, et al. RNAi screening reveals proteasome- and Cullin3-dependent stages in vaccinia virus infection. Cell Rep. 2012;2(4):1036–47.

57. Satheshkumar PS, Anton LC, Sanz P, Moss B. Inhibition of the ubiquitin-proteasome system prevents vaccinia virus DNA replication and expression of intermediate and late genes. J Virol. 2009;83(6):2469–79.

58. Teale A, Campbell S, Van Buuren N, Magee WC, Watmough K, Couturier B, et al. Orthopoxviruses require a functional ubiquitin-proteasome system for productive replication. J Virol. 2009;83(5):2099–108.

59. Moss B. Inhibition of HeLa cell protein synthesis by the vaccinia virion. J Virol. 1968;2(10):1028–37.

60. Parrish S, Moss B. Characterization of a vaccinia virus mutant with a deletion of the D10R gene encoding a putative negative regulator of gene expression. J Virol. 2006;80(2):553–61.

61. Parrish S, Moss B. Characterization of a second vaccinia virus mRNA-decapping enzyme conserved in poxviruses. J Virol. 2007;81(23):12973–8.

62. Rice AP, Roberts BE. Vaccinia virus induces cellular mRNA degradation. J Virol. 1983;47(3):529–39.

63. Strnadova P, Ren H, Valentine R, Mazzon M, Sweeney TR, Brierley I, et al. Inhibition of translation initiation by protein 169: a vaccinia virus strategy to suppress innate and adaptive immunity and alter virus virulence. PLoS Pathog. 2015;11(9):e1005151.

64. Weekes MP, Tan SY, Poole E, Talbot S, Antrobus R, Smith DL, et al. Latency-associated degradation of the MRP1 drug transporter during latent human cytomegalovirus infection. Science. 2013;340(6129):199–202.

65. Hunter S, Jones P, Mitchell A, Apweiler R, Attwood TK, Bateman A, et al. InterPro in 2011: new developments in the family and domain prediction database. Nucleic Acids Res. 2012;40(Database issue):D306–12.

66. Yu G, Luo H, Wu Y, Wu J. Ephrin B2 induces T cell costimulation. J Immunol. 2003;171(1):106–14.

67. Nakano K, Asano R, Tsumoto K, Kwon H, Goins WF, Kumagai I, et al. Herpes simplex virus targeting to the EGF receptor by a gD-specific soluble bridging molecule. Mol Ther. 2005;11(4):617–26.

68. Jafferji I, Bain M, King C, Sinclair JH. Inhibition of epidermal growth factor receptor (EGFR) expression by human cytomegalovirus correlates with an increase in the expression and binding of Wilms’ Tumour 1 protein to the EGFR promoter. J Gen Virol. 2009;90(Pt 7):1569–74.

69. Buller RM, Chakrabarti S, Cooper JA, Twardzik DR, Moss B. Deletion of the vaccinia virus growth factor gene reduces virus virulence. J Virol. 1988;62(3):866–74.

70. Henriksen L, Grandal MV, Knudsen SL, van Deurs B, Grovdal LM. Internalization mechanisms of the epidermal growth factor receptor after activation with different ligands. PLoS One. 2013;8(3):e58148.

71. Wiersma VR, Michalak M, Abdullah TM, Bremer E, Eggleton P. Mechanisms of Translocation of ER Chaperones to the Cell Surface and Immunomodulatory Roles in Cancer and Autoimmunity. Front Oncol. 2015;5:7.

72. Yue X, Wang H, Zhao F, Liu S, Wu J, Ren W, et al. Hepatitis B virus-induced calreticulin protein is involved in IFN resistance. J Immunol. 2012;189(1):279–86.

73. Cui X, Rouhani FN, Hawari F, Levine SJ. Shedding of the type II IL-1 decoy receptor requires a multifunctional aminopeptidase, aminopeptidase regulator of TNF receptor type 1 shedding. J Immunol. 2003;171(12):6814–9.

74. Cui X, Rouhani FN, Hawari F, Levine SJ. An aminopeptidase, ARTS-1, is required for interleukin-6 receptor shedding. J Biol Chem. 2003;278(31):28677–85.

75. Lin W, Zhang J, Lin H, Li Z, Sun X, Xin D, et al. Syndecan-4 negatively regulates antiviral signalling by mediating RIG-I deubiquitination via CYLD. Nat Commun. 2016;7:11848.

76. Smith GL, Talbot-Cooper C, Lu Y. How does vaccinia virus interfere with interferon? Adv Virus Res. 2018;100:355–78.

77. Talbot-Cooper C, Pantelejevs T, Shannon JP, Cherry CR, Au MT, Hyvönen M, et al. A strategy to supress STAT1 signalling conserved in pathogenic poxviruses and paramyxoviruses. BioRxiv. 2021.

78. Snary D, Barnstable CJ, Bodmer WF, Crumpton MJ. Molecular structure of human histocompatibility antigens: the HLA-C series. Eur J Immunol. 1977;7(8):580–5.

79. Parham P, Guethlein LA. Genetics of natural killer cells in human health, disease, and survival. Annu Rev Immunol. 2018;36:519–48.

80. Esteso G, Guerra S, Vales-Gomez M, Reyburn HT. Innate immune recognition of double-stranded RNA triggers increased expression of NKG2D ligands after virus infection. J Biol Chem. 2017;292(50):20472–80.

81. Garg A, Barnes PF, Porgador A, Roy S, Wu S, Nanda JS, et al. Vimentin expressed on Mycobacterium tuberculosis-infected human monocytes is involved in binding to the NKp46 receptor. J Immunol. 2006;177(9):6192–8.

82. Belin MT, Boulanger P. Processing of vimentin occurs during the early stages of adenovirus infection. J Virol. 1987;61(8):2559–66.

83. Risco C, Rodriguez JR, Lopez-Iglesias C, Carrascosa JL, Esteban M, Rodriguez D. Endoplasmic reticulum-Golgi intermediate compartment membranes and vimentin filaments participate in vaccinia virus assembly. J Virol. 2002;76(4):1839–55.

84. Aicher A, Hayden-Ledbetter M, Brady WA, Pezzutto A, Richter G, Magaletti D, et al. Characterization of human inducible costimulator ligand expression and function. J Immunol. 2000;164(9):4689–96.

85. Ogasawara K, Yoshinaga SK, Lanier LL. Inducible costimulator costimulates cytotoxic activity and IFN-gamma production in activated murine NK cells. J Immunol. 2002;169(7):3676–85.

86. Shiao SL, McNiff JM, Pober JS. Memory T cells and their costimulators in human allograft injury. J Immunol. 2005;175(8):4886–96.

87. Wallin JJ, Liang L, Bakardjiev A, Sha WC. Enhancement of CD8+ T cell responses by ICOS/B7h costimulation. J Immunol. 2001;167(1):132–9.

88. Sekiya T, Takaki S. RGMB enhances the suppressive activity of the monomeric secreted form of CTLA-4. Sci Rep. 2019;9(1):6984.

89. Xiao Y, Yu S, Zhu B, Bedoret D, Bu X, Francisco LM, et al. RGMb is a novel binding partner for PD-L2 and its engagement with PD-L2 promotes respiratory tolerance. J Exp Med. 2014;211(5):943–59.

90. Lorenzo MM, Sanchez-Puig JM, Blasco R. Mutagenesis of the palmitoylation site in vaccinia virus envelope glycoprotein B5. J Gen Virol. 2012;93(Pt 4):733–43.

91. Oie M, Shida H, Ichihashi Y. The function of the vaccinia hemagglutinin in the proteolytic activation of infectivity. Virology. 1990;176(2):494–504.

92. Chung CS, Chen CH, Ho MY, Huang CY, Liao CL, Chang W. Vaccinia virus proteome: identification of proteins in vaccinia virus intracellular mature virion particles. J Virol. 2006;80(5):2127–40.

93. Yoder JD, Chen TS, Gagnier CR, Vemulapalli S, Maier CS, Hruby DE. Pox proteomics: mass spectrometry analysis and identification of Vaccinia virion proteins. Virol J. 2006;3:10.

94. Resch W, Hixson KK, Moore RJ, Lipton MS, Moss B. Protein composition of the vaccinia virus mature virion. Virology. 2007;358(1):233–47.

95. McCraith S, Holtzman T, Moss B, Fields S. Genome-wide analysis of vaccinia virus protein-protein interactions. Proc Natl Acad Sci U S A. 2000;97(9):4879–84.

96. McSharry BP, Jones CJ, Skinner JW, Kipling D, Wilkinson GWG. Human telomerase reverse transcriptase-immortalized MRC-5 and HCA2 human fibroblasts are fully permissive for human cytomegalovirus. J Gen Virol. 2001;82(Pt 4):855–63.

97. McAlister GC, Huttlin EL, Haas W, Ting L, Jedrychowski MP, Rogers JC, et al. Increasing the multiplexing capacity of TMTs using reporter ion isotopologues with isobaric masses. Anal Chem. 2012;84(17):7469–78.

98. McAlister GC, Nusinow DP, Jedrychowski MP, Wuhr M, Huttlin EL, Erickson BK, et al. MultiNotch MS3 enables accurate, sensitive, and multiplexed detection of differential expression across cancer cell line proteomes. Anal Chem. 2014;86(14):7150–8.

99. Elias JE, Gygi SP. Target-decoy search strategy for increased confidence in large-scale protein identifications by mass spectrometry. Nat Methods. 2007;4(3):207–14.

100. Elias JE, Gygi SP. Target-decoy search strategy for mass spectrometry-based proteomics. Methods Mol Biol. 2010;604:55–71.

101. Kim W, Bennett EJ, Huttlin EL, Guo A, Li J, Possemato A, et al. Systematic and quantitative assessment of the ubiquitin-modified proteome. Mol Cell. 2011;44(2):325–40.

102. Wu R, Dephoure N, Haas W, Huttlin EL, Zhai B, Sowa ME, et al. Correct interpretation of comprehensive phosphorylation dynamics requires normalization by protein expression changes. Mol Cell Proteomics. 2011;10(8):M111 009654.

103. Huttlin EL, Jedrychowski MP, Elias JE, Goswami T, Rad R, Beausoleil SA, et al. A tissue-specific atlas of mouse protein phosphorylation and expression. Cell. 2010;143(7):1174–89.

104. Kall L, Canterbury JD, Weston J, Noble WS, MacCoss MJ. Semi-supervised learning for peptide identification from shotgun proteomics datasets. Nat Methods. 2007;4(11):923–5.

105. Makarov A, Denisov E. Dynamics of ions of intact proteins in the Orbitrap mass analyzer. J Am Soc Mass Spectrom. 2009;20(8):1486–95.

106. Ting L, Rad R, Gygi SP, Haas W. MS3 eliminates ratio distortion in isobaric multiplexed quantitative proteomics. Nat Methods. 2011;8(11):937–40.

107. van der Ploeg K, Chang C, Ivarsson MA, Moffett A, Wills MR, Trowsdale J. Modulation of Human Leukocyte Antigen-C by Human Cytomegalovirus Stimulates KIR2DS1 Recognition by Natural Killer Cells. Front Immunol. 2017;8:298.

108. Parkinson JE, Smith GL. Vaccinia virus gene A36R encodes a M(r) 43-50 K protein on the surface of extracellular enveloped virus. Virology. 1994;204(1):376–90.

109. Harris CR, Millman KJ, van der Walt SJ, Gommers R, Virtanen P, Cournapeau D, et al. Array programming with NumPy. Nature. 2020;585(7825):357–62.

110. McKinney W, editor Data structures for statistical computing in Python. 9th Pythn in Science Conference (SciPy, 2010); 2010.

111. Hunter JD. Matplotlib: A 2D graphics environment. Comput Sci Eng. 2007;9(3):90–5.

112. Virtanen P, Gommers R, Oliphant TE, Haberland M, Reddy T, Cournapeau D, et al. SciPy 1.0: fundamental algorithms for scientific computing in Python. Nat Methods. 2020;17(3):261–72.

113. Pedregosa F, Varoquaux G, Gramfort A, Michel V, Thirion B, Grisel O, et al. Scikit-learn: Machine Learning in Python. J Mach Learn Res. 2011;12:2825–30.

